# Extracellular vesicles induce aggressive phenotype of luminal breast cancer cells by PKM2 phosphorylation

**DOI:** 10.1101/2021.02.08.430192

**Authors:** Seo Young Kang, Eun Ji Lee, Jung Woo Byun, Dohyun Han, Yoori Choi, Do Won Hwang, Dong Soo Lee

**Affiliations:** Department of Molecular Medicine and Biopharmaceutical Science, Graduate School of Convergence Science and Technology, Seoul National University, Seoul, Korea; Department of Nuclear Medicine, Ewha medical center, Ewha Womans University College of Medicine, Seoul, Korea; Department of Nuclear Medicine, Seoul National University Hospital, Seoul National University College of Medicine, Seoul, Korea; Proteomics core facility, Biomedical Research Institute, Seoul National University Hospital, Seoul, Korea

**Keywords:** breast cancer cell, extracellular vesicles, aerobic glycolysis, PKM2 phosphorylation

## Abstract

The aerobic glycolysis is a hallmark of cancer glucose metabolism. Several studies have suggested that cancer-derived extracellular vesicles (EVs) can modulate glucose metabolism in adjacent cells and promote disease progression. Here we suggest that EVs originated from cancer cell with highly glycolytic activity can modulate glucose metabolism in the recipient cancer cells with relative low glycolytic activity, and further induce cell proliferation. Two types of breast cancer cell lines with different levels of glycolytic activity, MDA-MB-231 of a claudin low-type breast cancer cell and MCF7 of luminal type breast cancer cell, were selected and co-cultured using indirect co-culture system such as transwell system or microfluidic system. Glucose uptake of the recipient MCF7 cells was markedly increased after co-culture with MDA-MB-231 cells. MCF7 cells after co-culture with MDA-MB-231-tdTomato cells represented multiple tdTomato signal inside the cell, which proved that EVs originated from MDA-MB-231-tdTomato were transferred to MCF7 cell. In addition, serine phosphorylation of PKM2 necessary for tumorigenesis was highly activated, and tyrosine phosphorylation of PKM2 suggesting activated aerobic glycolysis was also increased in the co-cultured MCF7 cells. Proteomic profiling of the co-cultured MCF7 cells revealed the proliferation and dedifferentiation of MCF7 cells, and further confirmed epithelial-mesenchymal transition (EMT), which is a key phenomenon for cancer metastasis. In the transcriptomic analysis, glycolysis increased in co-cultured MCF7 cells, and the component analysis of genes associated with glycolysis revealed that the next major component after cytoplasm was extracellular exosome. Proteomic analysis of EVs revealed that there were important proteins in the EV such as EGFR, ERBB2 and MAPK for phosphorylating PKM2. This phenomenon suggests the potential for aggressive cancer cells to affect other cancer cells through EV mediators.

## Introduction

Cell-to-cell communication has well known to play a critical role during tumor progression and metastasis, allowing cancer cell to reprogram the surrounding tumor microenvironment (TME) (1, 2). The active crosstalk between cancer cells and tumor stroma consisting of immune and stroma cells, fosters various necessary processes for tumor progression and dissemination, including angiogenesis, cellular proliferation, immune-escape, formation of a pre-metastatic niche, invasion and multi-drug resistance (3, 4). Cell communication can be made through direct cell-to-cell interactions or the paracrine effect by soluble factors, such as cytokines, growth factors and chemokines (5). The most noteworthy way of communication between cancer cells and TME is the use of extracellular vesicle (EV) (6-8).

As a mechanism to communicate with the tumor microenvironment, tumor cells actively release large quantity of extracellular vesicles (EVs) that are a heterogeneous group of cell-derived membranous structures comprising exosomes and microvesicles (MVs) (9, 10). Exosomes are endosome-originated 50-150 nm small vesicles. MVs generated by the outward budding and fission of the plasma membrane have various size ranging from 50 nm to 1,000 nm in diameter (11, 12). These cancer cell-derived EVs, which are abundant in the body fluids of cancer patients, play a critical role in promoting tumor growth and progression through intercellular signaling (13, 14). Recently, several articles have been reported on the diagnosis and treatment of various diseases, and in particular on the role of EVs as therapeutic targets (15-18).

There is a study reporting on the potential of cancer-derived EVs modifying glucose utilization of recipient cells (2). The result suggests that miR-122 which is transferred to the normal cells through the EV inhibits the pyruvate kinase M2 (PKM2) of the recipient cells and lowers the glucose transporter 1 (GLUT1) to limit the glucose utilization of the recipient cells. It was also suggested in several studies that cancer cell-derived exosomes foster the tumor-like phenotype transform of other cells by activating glycolysis (19-21).

One thing to consider when we discuss activated glycolysis in cancer is the aerobic glycolysis, a characteristic glucose metabolism process of cancer. Aerobic glycolysis is a phenomenon observed by Otto Warburg that despite the availability of oxygen, most cancer cells produce energy predominantly by a high rate of glycolysis followed by lactic acid fermentation in the cytosol rather than by a comparatively low rate of glycolysis followed by pyruvate oxidation in the mitochondria (22-24). And the most important key molecule controlling aerobic glycolysis is pyruvate kinase (PK), which is precisely M2 isoform of pyruvate kinase (PKM2) (25-27).

Pyruvate kinase is the final rate-limiting protein of glycolysis and catalyzes the conversion of phosphoenolpyruvate (PEP) to pyruvate (28). There are multiple pyruvate kinase isoforms including PKM1, PKM2, PKL, and PKR expressed in different types of mammalian cells and tissues. The PKM2, resulting from a specific spliced mRNA form, mainly acts on cancer cells and is a key regulator of aerobic glycolysis known as the Warburg effect. Increased expression and phosphorylation of PKM2 is a characteristic glycolytic phenotype of cancer cells, which promotes rapid energy production and flow of glycolytic intermediates into collateral pathways to synthesize nucleic acids, amino acids, and lipids (28-30). Gambhir group, who recognized the value of PKM2 in cancer, developed positron emission tomography (PET) tracers, ^11^C-or ^18^F-labeled PKM2 specific radiotracer (DASA-23), for visualizing PKM2 *in vivo* (31-34). In addition, several studies have focused on PKM2 as a strategy to explore new anticancer drugs, and studies have shown that they may increase anticancer effect or reduce anticancer drug resistance. (35-37).

Although there are overwhelmingly many studies dealing with the interaction between cancer and stromal cells, studies have also been conducted to take into account the reciprocal interaction between cancer cells, albeit in a minority (38-40). As no one thinks a tumor consists of only one type of cancer cell, it is also worth considering the interactions between cancer cells of the tumor in terms of intratumor heterogeneity. Therefore, we investigate the role of EV modifying glucose metabolism in recipient cancer cells and major proteins inducing increased aerobic glycolysis, including PKM2, to figure out the reciprocal interactions between cancer cells.

## Materials and Methods

### Cell culture and analysis

#### Cell lines and cell culture

HepG2 (human hepatocellular carcinoma), Hep3B (human hepatocellular carcinoma), SK-OV-3 (human ovarian cancer cell), HT-1080 (human fibrosarcoma cell), HFF (human fibroblast), MDA-MB-231(human triple negative breast cancer cell) and MCF7 cells (human luminal type breast cancer cell) were purchased from Korean Cell Line Bank. The cells were cultured in Dulbecco’s modified Eagle’s medium (DMEM) containing with 10% fetal bovine serum (FBS), 10 U/ml penicillin, and 10 μg/ml streptomycin under humidified 5% CO_2_ atmosphere and at 37 °C.

MDA-MB-231 were transfected with a lentiviral vector encoding the red fluorescent protein palmitoylated-tdTomato (tdTomato) using lipofectamine transfection reagent (Life Technologies). Supernatant recovered after 48h from 293T transfected cells was filtered by a 0.45□μm pore membrane and added to MDA-MB-231 plated cells supplemented with 4□μg/mL polybrene (Sigma-Aldrich). After viral infection, palm-td-Tomato positive cells were selected using FACS for stable integration of the transfects.

#### Transwell cell culture

To induce indirect co-culture of two cell lines, transwell inserts with a PET membrane of 0.4-μm pore size (BD Bioscience) were used. MDA-MB-231 cells (0.8 × 10^5^ cells per well) were plated on the lower wells and MCF7 or HFF (1 × 10^4^ cells) were seeded into the top chamber (24-well insert; BD Bioscience) and then co-cultured for 24 hrs (Fig. 1a-b). All experiments were performed in triplicate.

**Figure 1.**
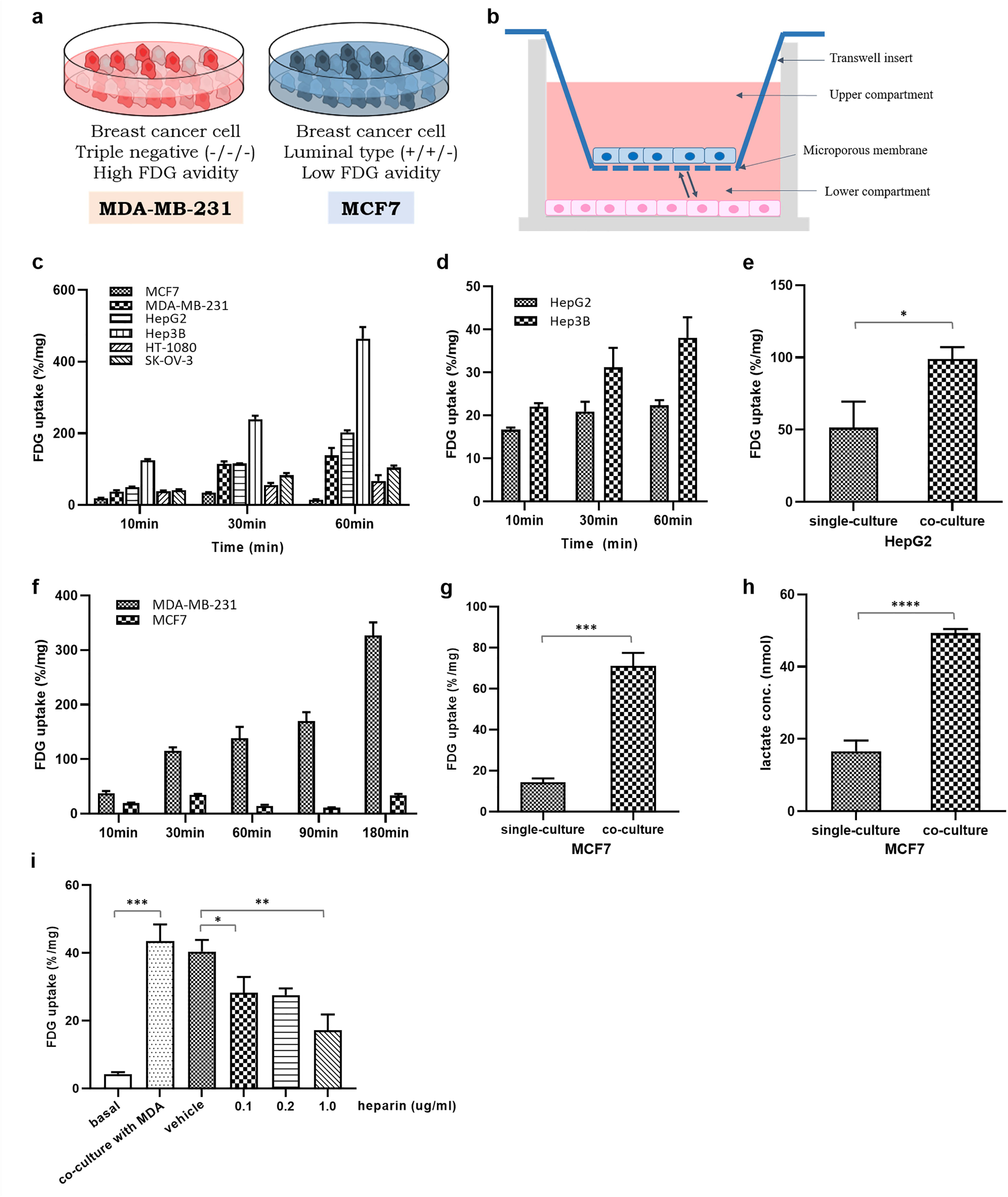
The change of glucose uptake in the MCF7 cell after co-culture with MDA-MB-231 cell. a, Two different breast cancer cell lines with different glycolytic activity. MDA-MB-231 and MCF7 cells; b, A schematic figure of a transwell system for indirect co-culture; c, Baseline FDG uptake of various cancer cell lines; d, Comparison of FDG uptake of HepG2 and Hep3B; e, The change of FDG uptake in the HepG2 after co-culture with Hep3B; f, baseline FDG uptake of two breast cancer cell lines; g, The change of FDG uptake in MCF7 cell after co-culture with MDA-MB-231; h, lactate assay in MCF7 cells after co-culture with MDA-MB-231; i, The change of FDG uptake increased by inhibiting EV uptake in MCF7 cells.

#### Glucose uptake analysis

Glucose analogue, ^18^F-fluorodeoxyglucose (FDG) was used to evaluate basal glucose metabolic status of these cells. After preculture for 4 hrs at glucose free media (glucose free DMEM, Gibco), cells were washed 3 times with PBS and cultured at 0.2 μCi of ^18^F-FDG with PBS for 1 hr at 37°C in 5% CO2. Cells were then harvested using 0.5% SDS solution and FDG uptake was measured using gamma counter (Packard Cobra-II Auto gamma counter). All experiments were performed in triplicate.

#### Heparin treatment

MCF7 cells pre-cultured with heparin at concentrations of 0.1, 0.2, and 1 ng/ml, respectively, were seeded on microfluidic chips with MDA-MB-231 cells. During the 24 hrs co-culture, the heparin concentration in the medium was maintained.

#### Lactate assay

The levels of lactate production were examined with a Lactate Assay Kit (DG-LAC200, DoGenBio, Korea). MDA-MB-231 cells (5 × 10^4^ cells per well) were plated in 24-well culture plate and the top chamber (24-well insert; BD Bioscience) were seeded with MCF7 (2 × 10^4^ cells) and then co-cultured for 24 hrs. After incubation for 24 hrs, lactate assays were performed with culture media collected from each sample according to the manufacturer’s protocol and the optical density was measured at 450 nm using a microplate reader (GloMax^®^-Multi E7031, Promega)

### Microfluidic system

#### Microfluidic Device Fabrication

A microfluidic device was established by the method mentioned in the previous report (41). It was manufactured by bonding Microchannel-patterned PDMS (polydimethylsiloxane; Sylgard 184; Dow Corning) to a glass coverslip. The conventional soft-lithography process was used to replicate the microchannel-patterned PDMS with SU-8 photoresist pattern master as a master mold (MicroChem). The PDMS elastomer, which was completely mixed with the curing agent at 10:1 weight ratio, was poured into the wafer and baked for 1 h 30 min in an oven at 80 °C. After curing, the PDMS replicas were removed from the wafer and all reservoir patterns on the PDMS replica were punched using skin biopsy punches. The sterilized PDMS replicas and glass coverslip were glued through oxygen plasma (Femto Science) and placed in the oven at 80 °C for at least 24 hrs to restore hydrophobicity of the microchannel surfaces.

#### Co-culture of MDA-MB-231 and MCF7 cells in the microfluidic system

Type 1 collagen ECM (2 mg/ml; BD Biosciences) was injected into the hydrogel channel and gelled in a 37°C incubator for 30 min. Cell seeding was then prepared by adding medium to the microfluidic channel and incubated at 37°C. MDA-MB-231 cells (donor cells, 200μL, 1 × 10^6^ cells/mL) were seeded in one reservoir of the cell culture channel (left channel) and MCF7 cells (recipient cell, 200μL, 1 × 10^6^ cells/mL) was seeded into the other reservoir of the cell culture channel (right channel). After cell attachment, in both reservoir of the culture channel, the medium was added and incubated at 37 °C in 5% CO_2_ atmosphere for 24 hrs.

### Preparation and characterization of Extracellular Vesicles (EV)

Extracellular vesicles from MDA-MB-231 were isolated using ultracentrifugation method. Filtered conditioned media was centrifuged in a Beckman Coulter Optima™ Ultracentrifuge at 150,000g at 4°C for 120 min with a Type 70 Ti rotor to pellet EVs. The supernatant was carefully removed, and crude EV-containing pellets were resuspended in ice-cold PBS and pooled. The amount of EV was estimated using the bicinchoninic acid assay (BCA; Thermo Scientific). Nanoparticle tracking analysis (NTA) system (NANOSIGHT NS500; Malvern) was used for measurement of EV size distribution and particle number. For optimal analysis, EVs were diluted with 1000-fold in particle-free PBS in the field of view. A laser beam of the system was adjusted to focus a suspension of the particles of interest. All measurements were recorded for further analysis by NTA software.

### Immunoassay

#### Immunoblotting

Cells were lysed in RIPA buffer, and centrifuged at 12,000 rpm for 30 min to remove cell debris. Protein concentration was measured using a BCA protein assay kit (Thermo Scientific). Equal amounts of protein per each sample were resolved by SDS-PAGE and transferred onto PVDF membranes (Millipore). The membrane was probed with rabbit monoclonal antibodies that recognize CD63 (1:1000; sc-5275; Santa Cruz biotechnology), CD81 (1:1000; sc-166029; Santa Cruz biotechnology), TSG101 (1:1000; ab83; Abcam), GLUT1 (1:1000; D3J3A; Cell Signaling), PKM2 (1:1000; D78A4; Cell Signaling), anti-phospho Y105 PKM2 (1:1000; ab156856; Abcam) overnight at 4 °C. Membranes were incubated with HRP-conjugated anti-rabbit secondary antibody for 2 h at room temperature, and proteins were visualized with a chemiluminescence detection system (Promega). Band intensities were quantified using AlphaView Software and results are expressed relative to the control condition. Three independent experiments were performed and the cropped version of the representative case of triplet is presented in the figures.

#### Immunocytochemistry

MCF7 cells co-cultured with MDA-MB-231 at transwell insert and microfluidic device were fixed with 4% paraformaldehyde (PFA) in PBS for 15 min at room temperature. The cells were then washed 3 times with PBS and treated with 0.5% Triton X-100 in PBS for 5 min at 4 °C to permeabilize cells. The samples were washed 3 times with PBS and non-specific binding sites were blocked with 5% BSA in PBS for 1 h at room temperature. Anti-rabbit GLUT1 (1:100; ab15309; Abcam), anti-rabbit PKM2 (1:100; D78A4; Cell Signaling), anti-rabbit phospho-PKM2 (1:100; PA5-78107; Thermo Fisher Scientific) antibodies were diluted in PBS and incubated with the samples at 4 °C overnight. In microfluidic system, diluted primary Abs were added into the microchannels and incubated at 4 °C overnight. The samples were then washed 3 times with PBS and incubated with Alexa Fluor 488-conjugated anti-rabbit secondary antibodies (1:10000; A32731; Invitrogen) for 1 h at room temperature to visualize the antibody reactions. Cell nuclei were stained using VECTASHIELD® mounting medium with DAPI (Vector Laboratories). Fluorescent images were acquired using a confocal laser scanning microscope (Leica TCS STED CW).

### Proteome profiling of cell lysates and extracellular vesicles

#### Cell culture and preparation of EV

Two cell lines, MDA-MB-231 and MCF7, cultured in DMEM medium containing 10% FBS, 100 μg/mL streptomycin and 100 U/mL penicillin using transwell inserts for in-direct col-culture. MDA-MB-231 cells (0.8 × 10^5^ cells per well) were plated on the lower wells of 24-well plate and the top chamber were seeded with MCF7 (1 × 10^4^ cells) and then co-cultured for 24 and 48 hrs. EV were isolated from MDA-MB-231 with same method as mentioned above.

#### Protein extraction

Proteins were extracted from the samples of both single-cultured and co-cultured MCF7 cells as well as EV originated from MDA-MB-231 cells in triplicate. The cell pellets were washed three times with cold PBS and lysed with 300 μl of lysis buffer (% SDS, 1 mM TCEP in 0.1 M TEAB pH 8.5). Protein concentration was measured using a BCA reducing agent compatible assay kit (Thermo Scientific, Rockford, IL, USA). Proteins were precipitated by adding ice-cold acetone overnight. Precipitated proteins were dissolved in 30 μl denaturation buffer (4% SDS and 100 mM DTT in 0.1M TEAB pH 8.5). After heating at 95□ for 15 min, denatured proteins were loaded onto 30 kDa amicon filter (Merck Millipore, Darmstadt, Germany). The buffer was exchanged with UA solution (8 M UREA in 0.1 M TEAB pH 8.5) by centrifugation at 14,000 g three times. After removal of SDS, cysteine alkylation was achieved by adding an alkylation buffer (50 mM IAA, 8 M UREA in 0.1 M Tris-HCl pH 8.5) for 1 hour at room temperature in the dark. Additional buffer exchanges were performed with 40mM TEAB pH 8.5 three times. The proteins were digested with trypsin (enzyme-to-substrate ratio [w/w] of 1:100) at 37□ overnight. The digested peptides were collected by centrifugation and the peptide concentrations were measured by tryptophan assay.

#### LC-MS/MS analysis and data processing for MCF7 cells

We distributed the 9 samples (3 groups of MCF7 cells with biological triplicate) to one TMT 10-plex set. Each 50 μg peptide sample was spiked with a uniform volume of ovalbumin. Subsequently, 40 mM TEAB buffer was added to each sample to equalize the volume. To eliminate errors caused by reagents, one set of TMT reagent was dissolved and spiked equally for the experiment. TMT reagent (0.8 mg) was dissolved in 110 μl of anhydrous ACN, of which 25 μl was added to the same channel in the experimental set. Then, 35 μl of ACN was added to achieve a final concentration of 30%. After incubation at room temperature for 1 hour, the reaction was quenched with hydroxylamine to a final concentration of 0.3% (v/v). The TMT-labeled samples were pooled at a ratio of 1:1 across all samples. The sample was lyophilized to almost dry and a desalting procedure was performed. The TMT-labeled peptide samples were desalted using solid phase extraction (SPE) column and high-pH peptide fractionation was performed by offline HPLC. Peptide samples were separated in a linear gradient and finally the sample was fractionated into 12 fractions. The fractions were lyophilized and stored at -80°C until MS analysis. The fractionated peptide samples were analyzed by LC-MS/MS system, a combination of an Easy-nLC 1000 (Thermo Fisher Scientific, Waltham, MA) connected to a nano electrospray ion source (Thermo Fisher Scientific, Waltham, MA) on Q-Exactive plus mass spectrometer (Thermo Fisher Scientific, Waltham, MA).

MS raw files were processed using Proteome Discoverer 2.1 software based on the SEQUEST-HT search engine against the Human Uniprot database. Database searches were performed using a 10-ppm precursor ion tolerance and a 0.02Da MS/MS ion tolerance. TMT tags on lysine residues and peptide N-termini and carbamidomethylation of cysteine residues were established as fixed modifications, while oxidation of methionine residues was set as a variable modification.

Statistical tests of the proteome were performed using Perseus software (42). After log2 transformation of the ion intensity, quantification was performed with a false discovery rate (FDR)-adjusted p-value < 0.05 with minimal fold-changes of ± 1.2. A comparative analysis between the single-cultured and co-cultured MCF7 cells identified total 57 differentially expressed proteins. There were 15 proteins identified in the first 24 hours, but eight more were found in the next 24 hours. Thirty-three more proteins were identified on a 48-hour incubation basis. Hierarchical clustering analysis revealed five clusters differentially expressed by co-culture duration among three groups. To functionally classify proteins of co-cultured MCF7 cells observed to be highly expressed relative to the single-cultured MCF7 cells in the proteomics data, we performed a gene ontology (GO) term analysis using Ingenuity Pathway Analysis (IPA, https://www.creative-proteomics.com/services/ipa-service.htm) (43) based on Uniprot database (http://www.uniprot.org/). Pathways were analyzed using the Kyoto Encyclopedia of Genes and Genomes (KEGG) databases. Protein-protein interactions (PPIs) for the network analysis were interrogated from STRING database (http://www.string-db.org) (44).

#### LC-MS/MS analysis and data processing for EVs from MDA-MB-231 cells

LC-MS/MS analyses were conducted with an Ultimate 3000 UHPLC system (Dionex, Sunnyvale, CA, USA) coupled to a Q-Exactive Plus mass spectrometer (Thermo Fisher Scientific, Waltham, MA). The column eluent was delivered to the Q-Exactive Plus via nanoelectrospray. In the data-dependent acquisition (DDA) method for label-free quantification, a survey scan (350 to 1650 m/z) was acquired with a resolution of 70,000 at m/z 200. A top-20 method was used to select the precursor ion with an isolation window of 1.2 m/z. The MS/MS spectrum was acquired at an HCD-normalized collision energy of 30 with a resolution of 17,500 at m/z 200. The maximum ion injection times for the full and MS/MS scans were 20 and 100 ms, respectively.

The raw LC-MS/MS data were analyzed using MaxQuant software (http://maxquant.org/, version 1.5.3.17) and searched against Human UniProt protein sequence database. Carbamido-methylation of cysteine was designated as a control variant, and N-acetylation of protein and oxidation of methionine were considered variable variants. Peptides with a minimum length of 6 amino acids and a maximum of 2 missing cuts were included. The acceptable false discovery rate (FDR) was set to 1% at the peptide, protein, and modification levels. For label-free quantification, an Intensity Based Absolute quantification (iBAQ) algorithm was used as part of the MaxQuant platform. The iBAQ values calculated by MaxQuant are the raw intensities divided by the number of theoretically observable peptides. Thus, the iBAQ values are provided in proportion to the molar amount of protein. Functional analysis of cargo proteins inside EVs was determined by GO enrichment of biological processes, molecular pathways and functions using the Functional Enrichment analysis tool (FunRich) version 3.1.3 (Funrich Industrial Co. Ltd, Hong Kong) (45, 46). The inter-protein pathways were analyzed using KEGG and the PPIs of network model were visualized using Cytoscape (47).

### Transcriptome profiling of cell lysates

#### RNA extraction

Total RNA were extracted from the cell lysates of both single-cultured and co-cultured MCF7 cells using Trizol reagent (Invitrogen). RNA quality was assessed by Agilent 2100 bioanalyzer (Agilent Technologies, Amstelveen, The Netherlands), and RNA quantification was performed using ND-2000 Spectrophotometer (Thermo Inc., DE, USA).

#### Library preparation and sequencing

Libraries were prepared from total RNA using the NEBNext Ultra II Directional RNA-Seq Kit (NEW ENGLAND BioLabs, Inc., UK). The isolation of mRNA was performed using the Poly(A) RNA Selection Kit (LEXOGEN, Inc., Austria). The isolated mRNAs were used for the cDNA synthesis and shearing, following manufacture’s instruction. Indexing was performed using the Illumina indexes 1-12. The enrichment step was carried out using of PCR. Subsequently, libraries were checked using the Agilent 2100 bioanalyzer (DNA High Sensitivity Kit) to evaluate the mean fragment size. Quantification was performed using the library quantification kit using a StepOne Real-Time PCR System (Life Technologies, Inc., USA). High-throughput sequencing was performed as paired-end 100 sequencing using HiSeq X10 (Illumina, Inc., USA).

#### Data analysis

A quality control of raw sequencing data was performed using FastQC (48). Adapter and low quality reads (<Q20) were removed using FASTX_Trimmer (49) and BBMap (50). Then the trimmed reads were mapped to the reference genome using TopHat (51). Gene expression levels were estimated using FPKM (Fragments Per kb per Million reads) values by Cufflinks (52). The FPKM values were normalized based on Quantile normalization method using EdgeR within R (53). Data mining and graphic visualization were performed using ExDEGA (E-biogen, Inc., Korea).

### Statistical analysis

All of the results were obtained from at least three independent experiments. Data are displayed as means ± standard deviation (SD). Comparison of results from experimental groups versus control groups were performed using R software (version 4.0.2, www.Rproject.org) and Prism (v8.0, GraphPad Software Inc.). A *p*-values of < 0.05.was considered statistically significant.

## Results

### Increased glycolytic activity of MCF7 after co-culture with MDA-MB-231 cells

Since human cancer cells have various glycolytic activity according to their aggressiveness, we compared glucose utilization status using glucose analogue, FDG, in several different cancer cell lines: MCF7, MDA-MB-231, HepG2, Hep3B, HT-1080, and SK-OV-3. As shown in Figure 1c, these cell lines showed various FDG uptake. To rule out the potential effects of high glucose media, we compared the effect of glucose uptake by the culture media. With DMEM including high glucose, MDA-MB-231 cells resulted in more increased FDG uptake, whereas Hep3B and HepG2 cells showed more increased FDG uptake with MEM including relatively low glucose. MCF7 cells did not represent significant difference. Although there is a difference in the FDG uptake by the culture medium, there is no difference in the overall tendency of the cellular uptake (supplementary Fig. S1a).

We selected two hepatoma cell lines, HepG2 and Hep3B to evaluate the change of FDG uptake according to co-culture between two cell lines. Hep3B is well known as aggressive hepatocellular carcinoma with highly glycolytic activity, whereas HepG2 has relatively low glycolytic activity (Fig. 1d). After co-culture with more aggressive cancer cell, Hep3B in transwell system, HepG2 showed increased FDG uptake compared to when cultured alone (Fig. 1e). We also compared the change of glycolytic activity in other cancer cell lines: luminal type breast cancer cell, MCF7 and triple negative breast cancer (TNBC) cell, MDA-MB-231. As we expected that TNBC cells are more aggressive and show increased glycolysis, MDA-MB-231 showed increased FDG uptake compared to MCF7 cells which have known as relatively less aggressive breast cancer cell (Fig. 1f). We examined the change of metabolic phenotypes such as glucose uptake and lactate production that represents an activated aerobic glycolysis in MCF7 cell after co-culture with MDA-MB-231 using transwell system. Again, MCF7 cells co-cultured with MDA-MB-231 for 24hrs showed dramatically increased FDG uptake compared to baseline level of MCF7 cell (Fig. 1g). More high level of lactate production suggesting activated aerobic glycolysis was also observed in co-cultured MCF7 cells compared to baseline of MCF7 cell (Fig. 1h). Since the main cause of this change was thought to be EVs secreted by Mda-MB-231, heparin, known to interfere with EV absorption in receiver cells, was used. The lowest heparin concentration tested, 0.1 μg/ml, achieved a 25 % reduction of FDG uptake in MCF7 cells co-cultured with MDA-MB-231 cells, and 50% reduced with 1ug/ml (Fig. 1i).

We also evaluated the effect of HepG2 and MCF7 cells on Hep3B and MDA-MB-231 cells. After co-culture with HepG2 and MCF7 cells in transwell system, respectively, FDG uptake of Hep3B and MDA-MB-231 cells did not significantly change compared to when cultured alone (supplementary Fig. S1b-c). While it is difficult to identify all the effects of relatively less aggressive cancer cells on more aggressive cancer cells, it has been confirmed that the glycolytic activity has not change much.

### Activated aerobic glycolysis of MCF7 cells via MDA-MB-231-derived EV

We postulated that EVs transported from MDA-MB-231 to the recipient MCF7 induced the change of glucose metabolism in the MCF7. First, we visualized MDA-MB-231-tdTomato cell-derived EVs inside MCF7 cells through co-culture using microfluidic chips mimicking interstitial fluid inside tumor distinct from macroscale transwell. Recipient MCF7 and donor MDA-MB-231-tdTomato cells were sequentially seeded 2-dimensionally in each channel and cultured for 24 hrs. Figure 2a showed that multiple red dot signals suggesting MDA-MB-231-tdTomato cell-derived EVs inside MCF7 cell and also in the channel between two cells of microfluidic chip (supplementary Fig. S2). Next, we isolated EVs from MDA-MB-231 cells using ultracentrifugation and nanoparticle-tracking analysis (NTA) of the isolated EVs showed a uniform size with the diameter of 80-150 nm (Fig. 2b). To verify the isolated EVs, expression of exosome-marker proteins, including CD63, CD61 and TSG101, was confirmed using western blot (Fig. 2c). FDG uptake of MCF7 treated with MDA-MB-231-derived EVs was significantly increased at concentration of 100 ug/ml (Fig. 2d) and CCK-8 assay showed increased cellular proliferation with EVs (Fig. 2e). To confirm whether MDA-MB-231-derived EVs regulates glucose metabolic phenotype in MCF7, we examined the change of FDG uptake in various culture condition of MCF7 such as EVs secreted from and EV-depleted conditioned media (EdCM) from MDA-MB-231 and HFF, respectively. MDA-MB-231-derived EV increased FDG uptake in MCF7 cells, although not as much as co-cultured. However, HFF-derived EVs did not induce a significant increase of FDG uptake. In addition, FDG uptake of MCF7 cultured in EdCM from two cell lines was slightly increased compared to baseline uptake of MCF7. Since the EdCM effects derived from the two cells did not show much difference, it seems likely to be non-specific effect caused by the growth factors secreted by the cells. (Fig. 2f).

**Figure 2.**
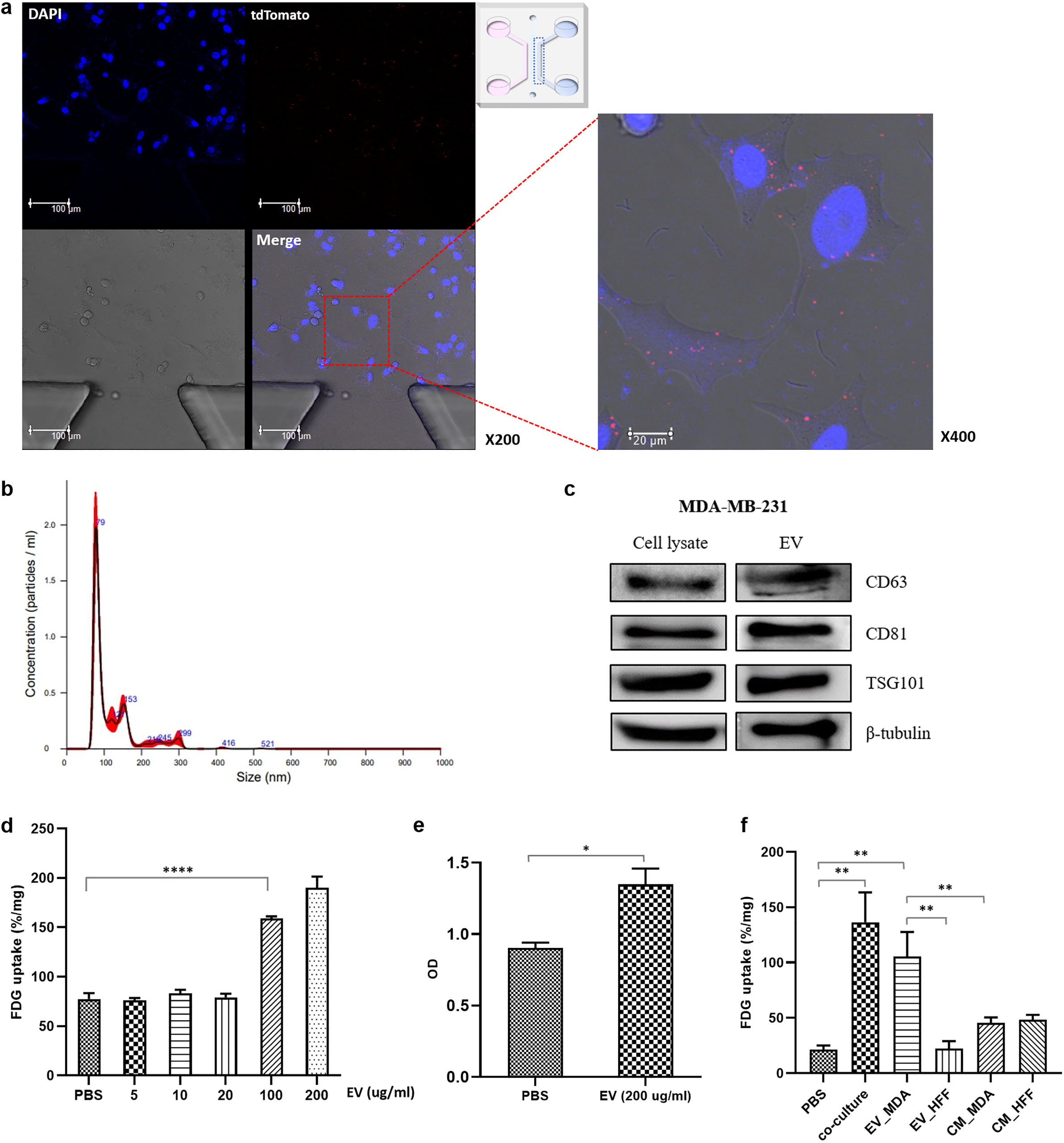
The impact of MDA-MB-231 cell-derived EVs for activation of glucose uptake in MCF7 cell. a, Confocal microscopic imaging shows tdTomato-EVs signals inside MCF7; b, nanoparticle tracking analysis (NTA) of isolated EVs from MDA-MB-231; c, Western blotting of EV-specific proteins, CD63, CD81, and TSG101; d, Increased dose-dependent FDG uptake of MCF7 after treatment with EVs isolated from MDA-MB-231; e, CCK8 assay showing increased cell proliferation of MCF7 after treatment with EVs isolated from MDA-MB-231; f, The effect of MDA-MB-231-mediated EVs compared to control groups such as EV-deprived conditioned medium and HFF-derived EVs.

When HFF was co-cultured with MDA-MB-231 cells, FDG uptake of HFF as well as MDA-MB-231 cells did not change significantly. MDA-MB-231-derived EVs increased FDG uptake at concentration of 200 ug/ml in HFF cells, but failed to induce proliferation of HFF cells. (supplementary Fig. S3).

### Tyrosine and serine phosphorylation of PKM2 for the regulation of glucose metabolism

We selected two representative proteins in glycolysis pathway: glucose transporter 1 (GLUT1) and pyruvate kinase muscle isozyme M2 (PKM2). GLUT1 is primarily responsible for the uptake of glucose into cancer cells. PKM2 is a rate-limiting glycolytic enzyme that catalyzes the final step in glycolysis, which is key in tumor metabolism and growth. On immunofluorescence analysis, depending on their function, GLUT1 was mainly distributed in the membrane and PKM2 in the cytosol. Since they are housekeeping proteins, it was difficult to identify changes in protein expression on microscopic imaging. (supplementary Fig. S4). However, serine phosphorylated PKM2 (S37) dramatically increased in co-cultured MCF7 cells compared to single cultured cells (Fig. 3a). Interestingly phosphorylated PKM2 signals were observed mainly in nucleus as well as cytosol of MCF7 cell. Several studies reported that serine phosphorylation of PKM2 (S37) induces its nuclear localization and performs non-glycolytic functions such as gene transcriptional regulation, which, in turn, facilitates metabolic reprogramming in cancer cells (54, 55).

**Figure 3.**
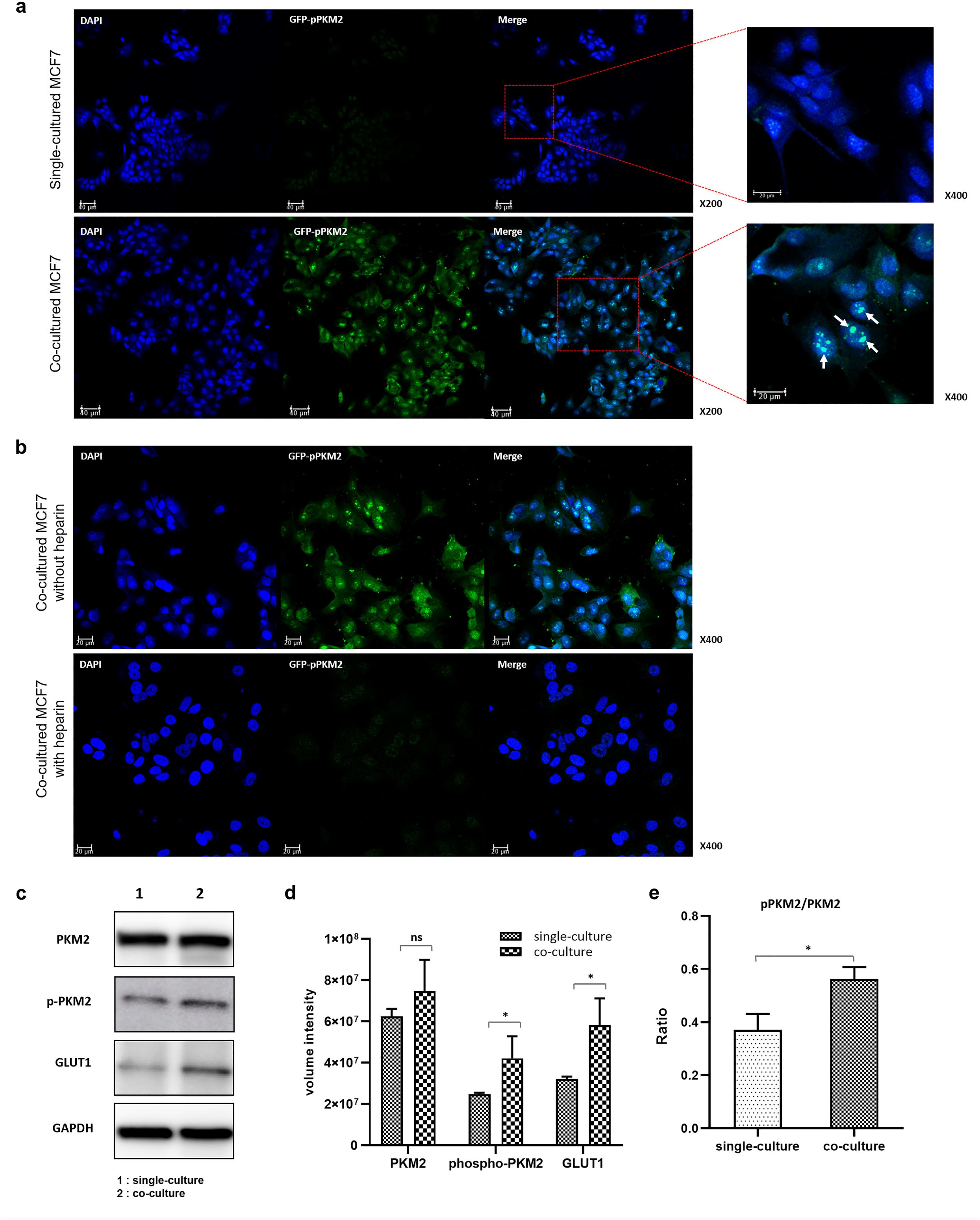
Increased phosphorylation of PKM2 in the MCF7 cell after co-culture with MDA-MB-231 cell. a, The phosphorylation of PKM2 S37 revealed a large increase in co-cultured MCF7 cells compared to single-cultured. Phosphorylated PKM2 is mainly located in nucleus. b, Suppressed activation of PKM2 phosphorylation by heparin, which inhibits EV uptake in the recipient cell, in the co-cultured MCF7 cell. The image also shows that the proliferation of cells has been inhibited. c, Western blotting of PKM2, phosphorylated PKM2, GLUT1 and GAPDH; d, Quantification of western blotting showed increased expression of GLUT1 and increased phosphorylation of PKM2 Y105, leading activation of aerobic glycolysis; e, Increased phosphorylation of PKM2 compared to expression of PKM2.

In our results, increased FDG uptake of MCF7 co-cultured with MDA-MB-231 cells was inhibited by heparin, known to interfere with EV uptake. Therefore, we assessed the change in expression of glycolytic pathway proteins by heparin treatment. Similarly, the expression of GLUT1 and PKM2 did not change much when determined visually. But even though the same amount of cells were seeded, the number of cells in the heparin treatment group showed a decrease, indicating that cell proliferation had decreased by inhibiting EVs uptake (supplementary Fig. S5). In addition, phosphorylation of PKM2 was greatly suppressed, showing a similar pattern to that seen in single-cultured MCF7 cells (Fig. 3b). That is, it can be considered that the effect of MDA-MB-231-derived EV disappeared due to inhibition of EV uptake. Therefore, it suggests that MDA-MB-231-derived EVs have upregulated aerobic glycolysis by activating the phosphorylation of PKM2 in MCF7 cells and consequently induced cell proliferation.

Western blot analyses showed that expression of GLUT1 and PKM2 were increased in the co-cultured MCF7 cell compared to single-cultured one, although not statistically significant in PKM2. Tyrosine phosphorylation of PKM2 (Y105) was significantly increased in the co-cultured MCF7 cells, again suggesting that aerobic glycolysis was stimulated by co-culture with MDA-MB-231 cells (Fig. 3c-d). Especially, ratio of PKM2 phosphorylation to PKM2 was significantly increased in the co-cultured MCF7 cells, suggesting phosphorylation of PKM2 was a key stimulant for activated aerobic glycolysis (Fig. 3e).

### Proteomic analysis of MCF7 co-cultured with MDA-MB-231 cells

Through the above experiments, we were able to confirm the role of EVs in regulating the glucose metabolism of recipient cells. For an overall analysis of the changes induced by MDA-MB-231 cell-derived EV, we analyzed the proteome profile of MCF7 cells co-cultured with MDA-MB-231 cells for 24h and 48h respectively, and single cultured as a control. ANOVA analysis of three groups identified five protein clusters according to expression pattern, two major clusters, 379 and 381, of them that had a consistent direction of expression were selected for further analysis (Fig. 4a). According to GO term analysis using IPA, the proteins contained in cluster 379 were mainly classified into the following categories: regulation of cell communication, nucleobase-containing compound biosynthetic process, cellular response to organic substance, positive regulation of gene expression and so on. On the other hand, proteins in cluster 381 mainly represented gene ontology associated with cell dedifferentiation, such as cell morphogenesis and regulation of cell differentiation (Table 1.).

**Table 1.**
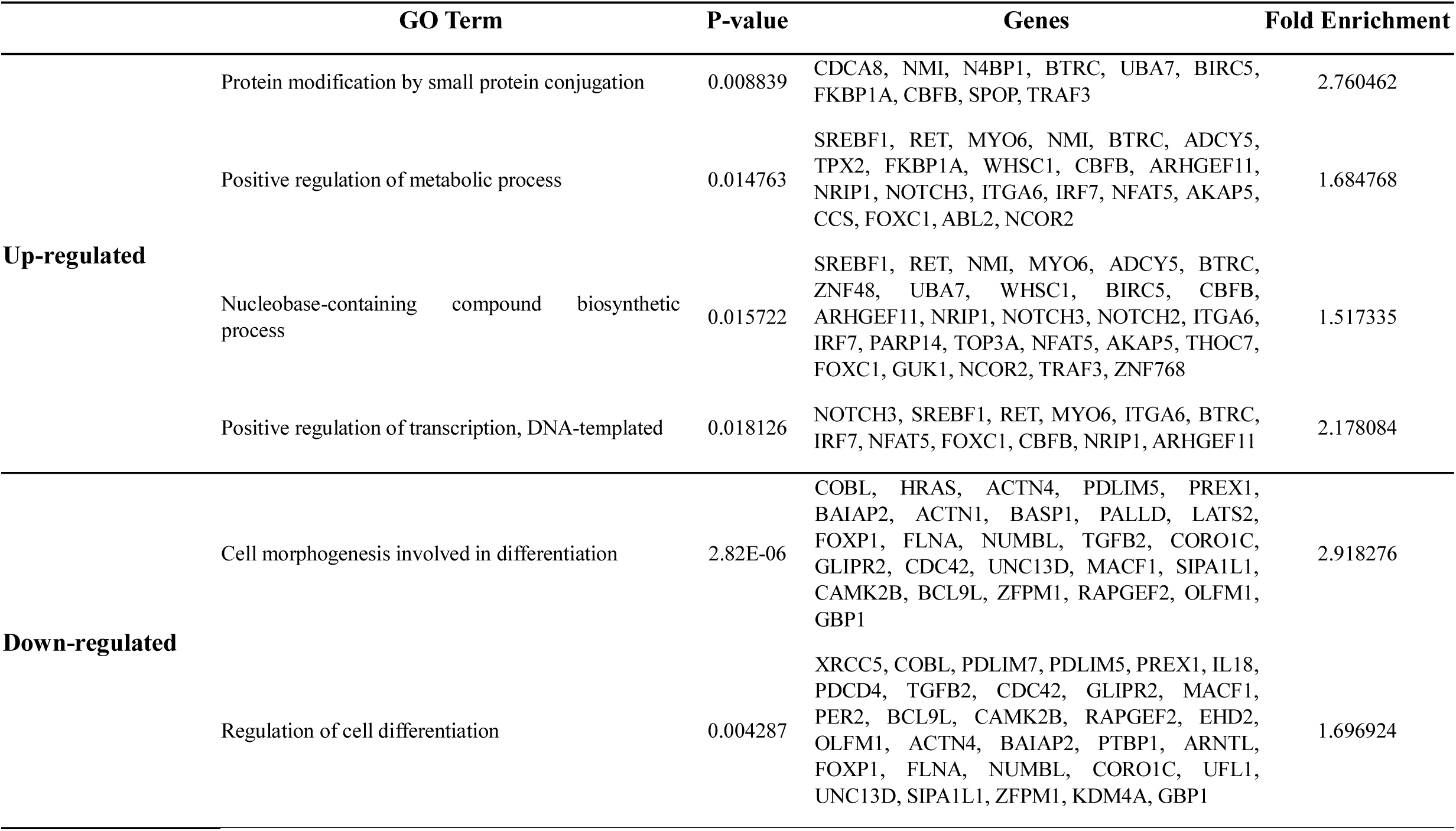
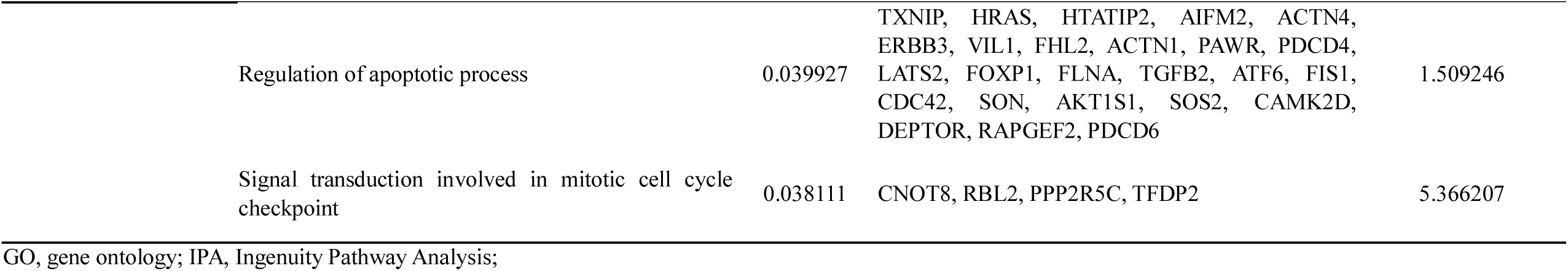
GO term analysis using IPA represent distinct protein groups related to biosynthesis and differentiation.

**Figure 4.**
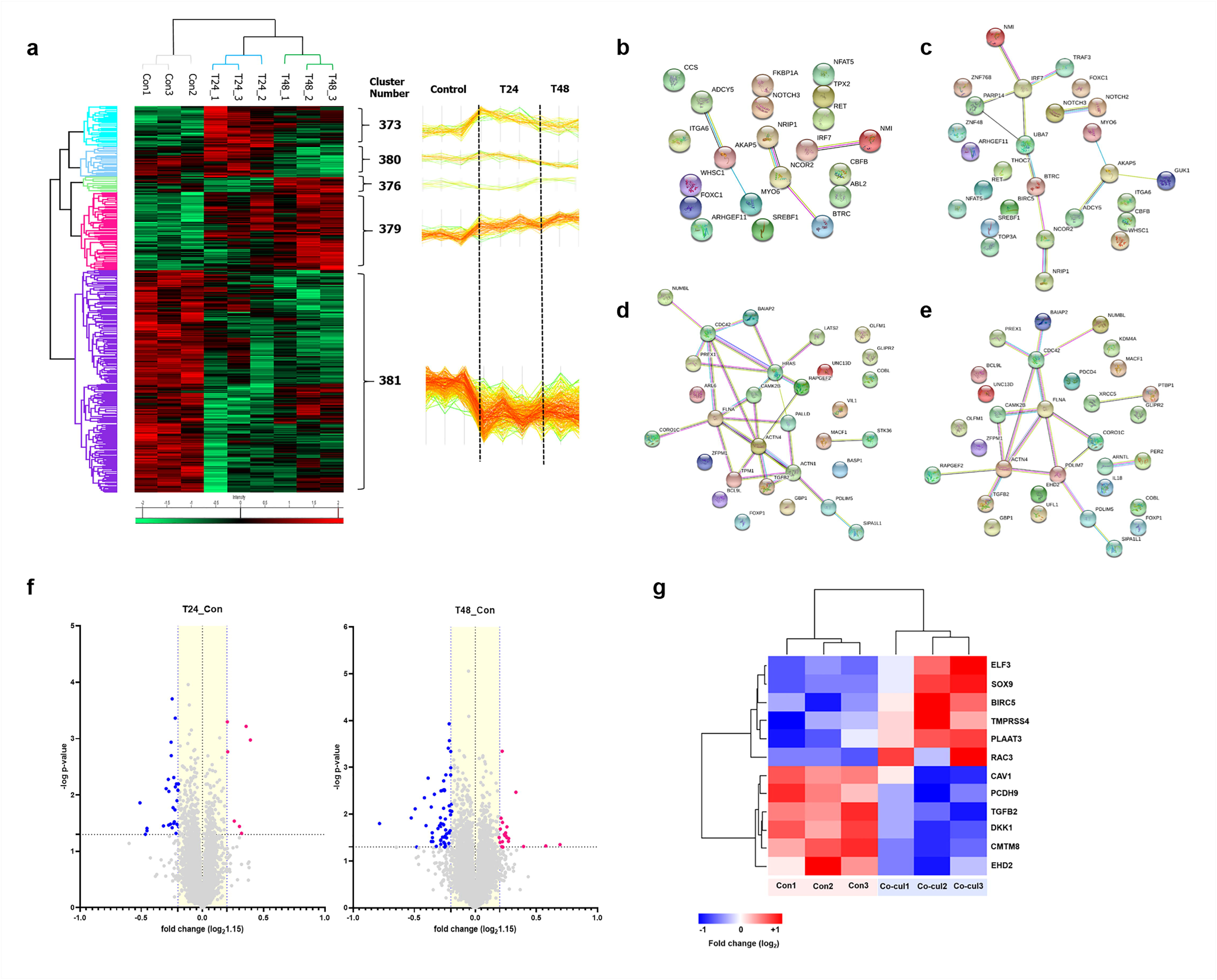
Proteomics analysis. a, Hierarchical clustering represented five clusters, and of them, we noted the major two cluster (#379 and #381) with consistent direction in expression. b, Protein-protein interactions of the differentially expressed proteins in MCF7 cells were visualized using STRING analysis. The clusters of up-regulation: b, positive regulation of metabolic process; c, nucleobase-containing compound biosynthetic process. The clusters of down-regulation: d, cell morphogenesis; e, regulation of cell differentiation. Lines represented interactions between proteins and thickness denoted the confidence level associated with each interaction. f, Volcano plotting described differentially expressed proteins in the MCF7 cell co-cultured with MDA-MB-231 (-log p-value>1.3 and fold change≥1.15). Thirty-two differentially expressed proteins after co-culture for 24h and 74 proteins for 48h were identified; d, Several proteins associated to EMT were expressed differentially in the MCF7 cells co-cultured for 48 hours compared to single-cultured cells.

Protein-protein interaction for differentially expression proteins was visualized using STRING analysis, and main interactive clusters, upregulated of nucleobase-containing compound biosynthetic process, and down-regulated cell morphogenesis and regulation of cell differentiation, were displayed (Fig. 4b-e). Comparative analysis of the single-cultured and co-cultured MCF7 cells identified 32 differentially expressed proteins for 24h and 74 proteins for 48h (Fig. 4f). Except for overlapping proteins, a total of 96 differentially expressed proteins were identified. Among them, markers of the epithelial phenotypes such as the tight junction protein ZO-1 (TJP1) and protocadherin beta-2 (PCDHB2) were reduced in MCF7 cells co-cultured with MDA-MB-231 for the first 24 hours. On the other hand, proteins such as ETS-related transcription factor (ELF3), transcription factor SOX-9 (SOX9), ras-related C3 botulinum toxin substrate 3 (RAC3), Caveolin-1 (CAV1), and Dickkopf-related protein 1 (DKK1), which were well known as EMT driver, were differentially expressed in co-cultured MCF7 cells for 48 hours (Fig. 4g).

### Transcriptomic analysis of MCF7 co-cultured with MDA-MB-231 cells

Transcriptomic analysis was performed to analyze the pattern of early expression changes at the RNA level of MCF7 cells co-cultured with MDA-MB-231 cells for 24h. Total 452 genes with more than 1.5 fold changes were differentially expressed in co-cultured MCF7 cells (Fig. 5a-b). According to GO analysis, they were mainly related to glucose metabolism, apoptosis, cell cycle, cell differentiation, and cell proliferation, especially the genes related to glucose metabolism were highly up-regulated (Fig. 5c-d). The results of biological processes showed that genes were highly related to reactions to hypoxic, nucleotide phosphorylation, purine metabolic process and glycogenesis. (Fig. 5e). In the analysis of cellular component, the main cellular component after cytosol and nucleus was the extracellular exosome (Fig. 5f). As a result of analyzing the cellular component of genes related to glycolysis in the entire transcriptomic data, the highest cellular component after cytosol was also the extracellular exosome (supplementary Fig. S6). This suggests that a number of glycolysis-related RNA genes inside MCF7 cells were carried by MDA-MB-231-derived EV. Significant changes in glycolysis/gluconeogenesis were also observed in the KEGG pathway analysis (Fig. 5g).

**Figure 5.**
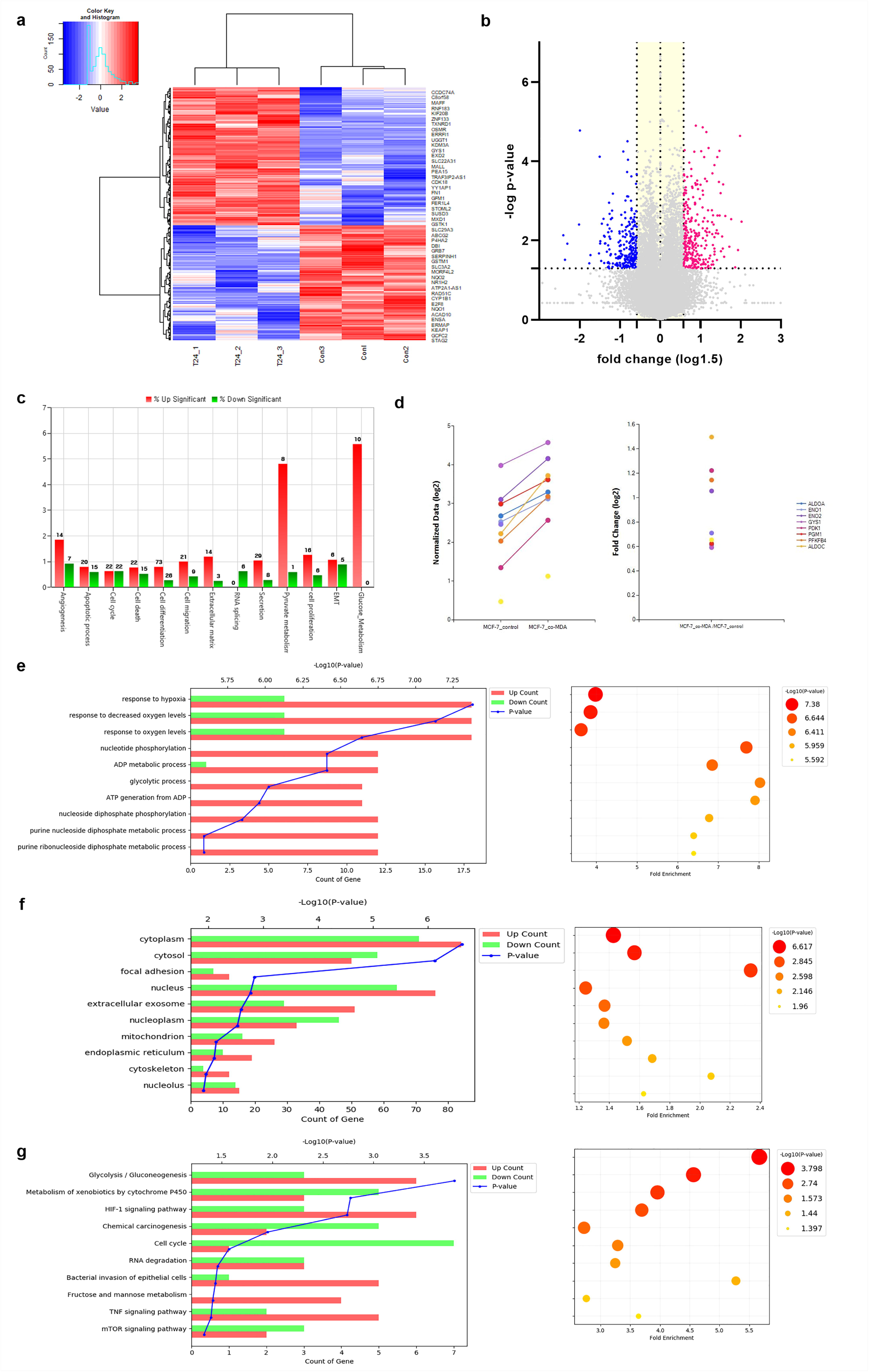
Transcriptomic analysis. a, Hierarchical clustering analysis of transcriptomic data showed differentially expressed mRNAs in the co-cultured MCF7 cells with MDA-MB-231 for 24h; b, Volcano plotting described 452 differentially expressed genes in the MCF7 cells co-cultured with MDA-MB-231 (-log p-value>1.3 and fold change≥1.5); c, Gene category analysis revealed that the differentially expressed genes were mainly related to glucose metabolism, apoptosis, cell cycle, cell differentiation, and cell proliferation. d, The genes related to glucose metabolism were highly up-regulated; e, The biological process analysis showed that they were highly related to response to hypoxia, nucleotide phosphorylation, purine metabolic process and glycolysis; f, The extracellular exosome component were shown to be significant in the cell component analysis. g, KEGG pathway analysis also showed significant changes in glycolysis/gluconeogenesis.

### Cargo proteins of MDA-MB-231-derived EVs

Next, we examined proteome profiles of EVs from MDA-MB-231 cells to find the key molecules that led to these changes, resulting in the identification of a total of 856 proteins in biological triplicate. We compared the 856 proteins with previously published data in Vesiclepedia databases. Our results indicated overlaps of 823 (96%) and 685 (80%) of proteins among total identified EV and MDA-MB-231 cell-derived EV in Vesiclepedia database, respectively (supplementary Fig. S7a). To investigate the biologic function of the identified proteins, GO analysis was conducted. Proteins were sorted into categories according to their ontology as determined from their GO annotation terms. The annotated biologic processes of the proteins revealed enrichment of EV-associated proteins related to RNA binding, translational initiation, and post-translational protein modification (supplementary Fig. S7b). In addition, analysis of the KEGG pathway of EV proteins revealed total 360 signal pathways, and there were 36 significant pathways consisting of a considerable number of proteins. Among them were glycolysis/gluconeogenesis, pyruvate metabolism, and PI3K-Akt signaling pathways (supplementary Fig. S8; supplementary Table S1). PI3K signaling pathway contains proteins such as EGFR, ERBB2, and MAPK1, which are well known for phosphorylating PKM2. Therefore, we suggest the possibility that the EGFR, ERBB2, and MAPK1, the cargo proteins of EV, have phosphorylated PKM2 and resulted in activated aerobic glycolysis and cell proliferation.

## Discussion

Cell-to-cell communication is an essential process for cell biology and function of in vivo, and many elements are organically connected to facilitate theses interactions. Among them, the most recent thing that has attracted great attention is the extracellular vesicles. Extracellular vesicles comprise a heterogeneous population of membrane vesicles of various origins. Their size may vary (typically between 50 nm and 500 nm, but they can be even larger, measuring 1-10 μm). Over the past two decades, extracellular vesicles have been named based on their origin (cell type), size, morphology and cargo content but can now be classified into two distinct classes: exosomes and microvesicles (9). There are a few articles that evaluated the role of EVs modulating glucose metabolism in recipient cells (2, 56). But, most studies focus primarily on the relationship between cancer and stromal cells, so there is little research on the communication of cancer cells through EVs. Recently, research on circulating tumor cells (CTC) or circulating tumor DNA (ctDNA) has become more active, suggesting the existence of various kinds of subclones in a tumor (57-59). In this respect, we investigated the interaction between cancer cells through EVs derived from cancer cells, and among them focused on aerobic glycolysis, which is a characteristic of cancer.

Our findings suggest that MDA-MB-231-derived EVs instigated phosphorylation of PKM2, resulted in increased aerobic glycolysis and cell proliferation, that is, transformed MCF7 cells more aggressively. These imply the possibility of interaction between subclones within a tumor in terms of cell-to-cell communication and tumor heterogeneity. There are a few studies dealing with the interaction between cancer cells in a tumor (38-40). Rak et al., reported that oncogenic receptor tyrosine kinase, EGFRvIII, were transferred to EGFR-negative endothelial cells (HUVECs) and A431 cells expressing only wtEGFR through microvesicles derived from EGFRvIII-positive glioblastoma cells, resulting in activation of downstream signaling pathway such as MAPK and Akt (37). In a similar way, in our result, MCF7 co-cultured with more aggressive subtype, MDA-MB-231 cells, revealed dramatically increased FDG uptake compared to baseline uptake level. EVs isolated from MDA-MB-231 cells also had a similar effect, although not as effective as co-cultured. Increased cellular proliferation and lactate production accompanied by increased FDG uptake are signified to be caused by activated aerobic glycolysis rather than oxidative phosphorylation. However, EVs derived from HFF did not arouse this phenomenon (Fig. 2f). Therefore, it can be inferred that these have been wrought by a special cargo of MDA-MB-231-derived EVs. EV-deprived conditioned media from MDA-MB-231 and HFF cells also produced a significant glucose uptake in the recipient MCF7 cells, although not as much as EVs, which is explained by the high volume of substances including growth factors secreted from cells. Given the significant difference between the effects of EV and CM, this does not mean that the effects of EV are meaningless.

It has been well established that tumor cells have elevated rates of glucose uptake and high lactate production in despite of the presence of oxygen, known as aerobic glycolysis (60). Glucose-metabolizing enzymes, including pyruvate kinase, are often upregulated in cancer cells (61, 62). Pyruvate kinase (PK) regulates the final rate-limiting step of glycolysis and catalyzes the transfer of a phosphate group from phosphoenolpyruvate (PEP) to adenosine diphosphate. There are four mammalian pyruvate kinase isoforms (PKM1, PKM2, PKR, PKL), and PKM2 is found predominantly in the proliferative cells such as fetus and cancer (29). PKM2 has two isoforms, tetrameric and dimeric forms, and exhibits a high level of catalysis when protein is in a tetrameric state but provides a growth advantage in a dimeric state (63). PKM2 convert to a dimeric form when Y105 is phosphorylated, and the dimeric form of PKM2 has low affinity for the precursor, PEP, leading to the aerobic glycolysis with increased lactate production (64). Unlike tyrosine phosphorylation of PKM2 that principally modulates its glycolytic function, serine phosphorylation of PKM2 supports its nuclear localization (54, 55, 65) and essential for tumorigenesis (66).

Confocal microscopic images stained with phospho-PKM2-S37 antibody described increased signals in nucleus with relatively week but clear uptake in cytosol of co-cultured MCF7 cells. To be exactly, the main signals seems to be located in nucleolus. The nucleolus is a distinct subnuclear compartment and primarily associated with ribosome biogenesis. However, several lines of evidence represent it has additional functions, such as regulation of mitosis, cell-cycle progression and proliferation (67). The S37-phosphorylated PKM2 assists the transcriptional activation of signal transducer and activator of transcription 5 (STAT5) to facilitate the expression of different genes such as cyclin D1, c-Myc, glucose transporter 1 (GLUT1), lactate dehydrogenase A (LDHA), and PKM2, all essential for tumor cell metabolic reprogramming and proliferation (29, 54, 55). Therefore, nuclear localization of S37 phosphorylated PKM2 stimulated in co-cultured MCF7 cells supports that MDA-MB-231-derived EVs lead conversion of MCF7 cells to more aggressive cancer cell. In addition, after adding heparin that interfered with the EV uptake of MCF7 cells, activation of S37 phosphorylation in co-cultured MCF7 cell was suppressed. Cell proliferation was also reduced in the heparin treated group. This once again reveals the role of MDA-MB-231-derived EVs, indicating the need to uncover the major substances inside the EV that caused this reaction. What’s interesting is that the S37 phosphorylation signal was feeble in a single cultured MCF7 cells (Fig. 3a), compared with very strong S37 phosphorylation in single-cultured MDA-MB-231 cells (supplementary Fig. S9). This is probably related to the less proliferative nature of MCF7 cells, and further research is needed.

MDA-MB-231 cells represented different patterns in expression of GLUT1 and phosphorylation of PKM2. Unlike MCF7 cells, most of the GLUT1 proteins was located in the cytosol like granules. It can be interpreted that the highly expressed GLUT1 proteins are in the endoplasmic reticulum or Golgi apparatus. In addition, S37 phosphorylation of PKM2 in MDA-MB-231 cells was highly activated and located throughout the cell as well as nucleus (supplementary Fig. S9). Although both are breast cancer cells, these differences between MCF7 and MDA-MB-231 cells seems to be due to a fundamental heterogeneous nature. However, by MDA-MB-231-derived-EVs, the pattern of PKM2 phosphorylation in MCF7 cells was changed similarly to MDA-MB-231 cells. Therefore, it can be a clue to the transfer of characteristic substances by EV between different cells.

Proteomics analysis results revealed that a total number of 97 proteins were differentially expressed in MCF7 co-cultured with MDA-MB-231 compared to single-cultured MCF7 cells. GO analysis showed that cell differentiation decreased, whereas gene expression and translation increased in the co-cultured MCF7 cells. This is in line with the results of FDG uptake test, immunoblot and immunostaining, which show increased glucose uptake and phosphorylation of PKM2. In addition, a number of EMT-related proteins existed among 97 differentially expressed proteins. Marker protein of epithelial phenotypes such as TJP1 and PCDHB2 began to decrease in expression in co-cultured MCF7 cell of 24 hours, and after co-culture of 48 hours, the expression of proteins such ELF3, SOX9, RAC3, DKK1, and CAV1, known as EMT driver, also increased or decreased according to their role. Considering the characteristics of the MDA-MB-231 cell of the claudin-low type, which is enriched in EMT features, it can be interpreted that the EV derived from MDA-MB-231 delivered these characteristics to the MCF7 cell. As mentioned above, nuclear PKM2 is well known to act as a transcription factor, contributing to the expression of several proteins involved in cell proliferation. Furthermore, it is also known to interact with TGF-β-induced factor homeobox 2 (TGIF2) and to repress E-cadherin expression during EMT, which plays a critical role in the transition to mesenchymal phenotype of cancer cell (68, 69). Therefore, the special cargo protein in EV not only activated aerobic glycolysis by phosphorylating PKM2 but also induced nuclear transfer of PKM2 to result in EMT occurring.

Transcriptomic analysis revealed that a total number of 456 genes were differentially expressed in MCF7 co-cultured with MDA-MB-231 compared to single-cultured MCF7 cells. GO analysis showed that differentially expressed genes (DEGs) were related glycolysis and cell proliferation, which is consistent with the results from FDG uptake test and proteomics data. Interestingly, extracellular exosome was the third highest after cytosol and nucleus for DEGs in the cell component analysis, and in the component analysis of glycolysis-related genes among the entire transcriptomic data, extracellular exosome was the second highest cell component after cytosol. These results suggest that several genes, especially those related to glycolysis, may have carried into MCF7 through EV. This is thought to be the main result to support this research hypothesis.

It was thought that the cause of this remarkable changes in MCF7 cells was the special cargo in the MDA-MB-231-derived EV. To confirm this, the proteomic analysis was implemented for MDA-MB-231-derived EVs. Total 856 proteins were identified and their analysis revealed a number of significant KEGG pathway. Among them were glycolysis/gluconeogenesis, pyruvate metabolism and PI3K-Akt signaling pathway. PI3K-Akt signaling pathway promotes metabolism, proliferation, cell survival, and growth and contains a number of proteins such as EGFR, ERBB2, and MAPK1, associated with their function. ERBB2, one of oncogenic tyrosine kinases that are frequently activated in breast cancers can phosphorylate PKM2 at Y105 (66). EGFR and MAPK1 are also famous oncogenic kinases and have the ability to phosphorylate PKM2 at S37 (50, 51). Therefore, we suggest the possibility that the EGFR, ERBB2, and MAPK1, the cargo proteins of EV, have phosphorylated PKM2 and resulted in activated aerobic glycolysis and cell proliferation, even transition to mesenchymal phenotype in the MCF7 cells co-cultured with MDA-MB-231 cells (Fig. 6). In addition, In addition, the presence of glycolysis pathway in the EV protein analysis is also an undeniable finding. This, too, would have led to the transmission of major proteins associated with glycolysis, which would have contributed to the activation of aerobic glycolysis in MCF7 cells.

**Figure 6.**
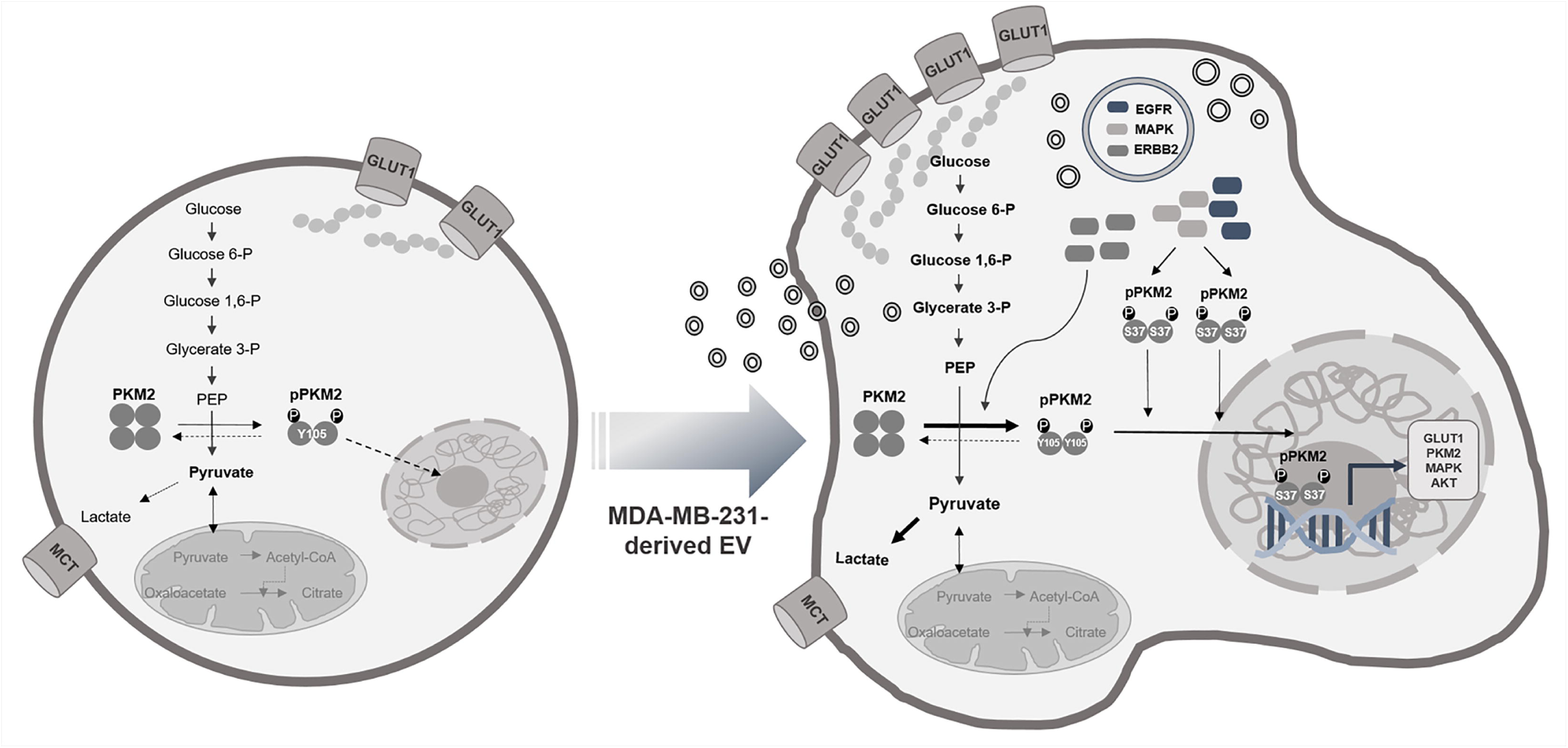
Extracellular vesicles as carriers for the transfer of aggressive phenotypes. The cargo proteins of EVs, EGFR, ERBB2, and MAPK1, induced phosphorylation of PKM2 and resulted in activated glycolysis and cell proliferation, even transition to mesenchymal phenotype of the MCF7 cell.

EVs originated from MDA-MB-231 cells have upregulated aerobic glycolysis by inducing phosphorylation of PKM2 in the MCF7 cells, which finally made dedifferentiation and cellular proliferation. Therefore, given the ability of EVs originated from MDA-MB-231 cells to cause tumor progression of other cancer cells, we note the potential impact of aggressive cancer cells on other cancer cells nearby in terms of intratumor heterogeneity. This impact may be caused by an intermediary called EVs.

