## Supplemental material for "Extracellular vesicles induce aggressive phenotype of luminal breast cancer cells by PKM2 phosphorylation"

**Supplementary** **Table S1**. KEGG pathway analysis from proteomics of MDA-MB-231-derived EVs

| **KEGG Pathway** | **P value** | **Genes** | **% associated genes** |
| --- | --- | --- | --- |
| PI3K-Akt signaling  pathway | 0.02540566 | COL1A1, COL1A2, COL2A1, COL4A1, COL4A2, COL4A5, COL6A1, COL6A2, COL6A3, EGFR, EPHA2, ERBB2, ERBB4, FN1, GNB1, GNB2, GNB3, GNB4, GNG12, GNG5, HRAS, HSP90AA1, HSP90AB1, IGF2, ITGA2, ITGA2B, ITGA3, ITGA6, ITGAV, ITGB1, ITGB3, KRAS, LAMA3, LAMA5, LAMB1, LAMB2, LAMB4, LAMC1, MAPK1, NRAS, PDGFC, RAC1, RELN, RPS6, THBS1, THBS2, THBS3, THBS4, TNC, VWF, YWHAB, YWHAE, YWHAG, YWHAH, YWHAQ, YWHAZ | 12.99% |
| Glycolysis/Gluconeogenesis | 0.00103427 | ADH5, ALDH7A1, ALDH9A1, ALDOA, ALDOC, BPGM, ENO1, ENO2, ENO3, GAPDH, GAPDHS, GPI, LDHA, LDHAL6A, LDHB, LDHC, PFKL, PFKM, PFKP, PGAM1, PGAM2, PGK1, PGK2, PKLR, PKM, TPI1 | 25.0% |

KEGG, Kyoto Encyclopedia of Genes and Genomes;

**Supplementary Figure S1**

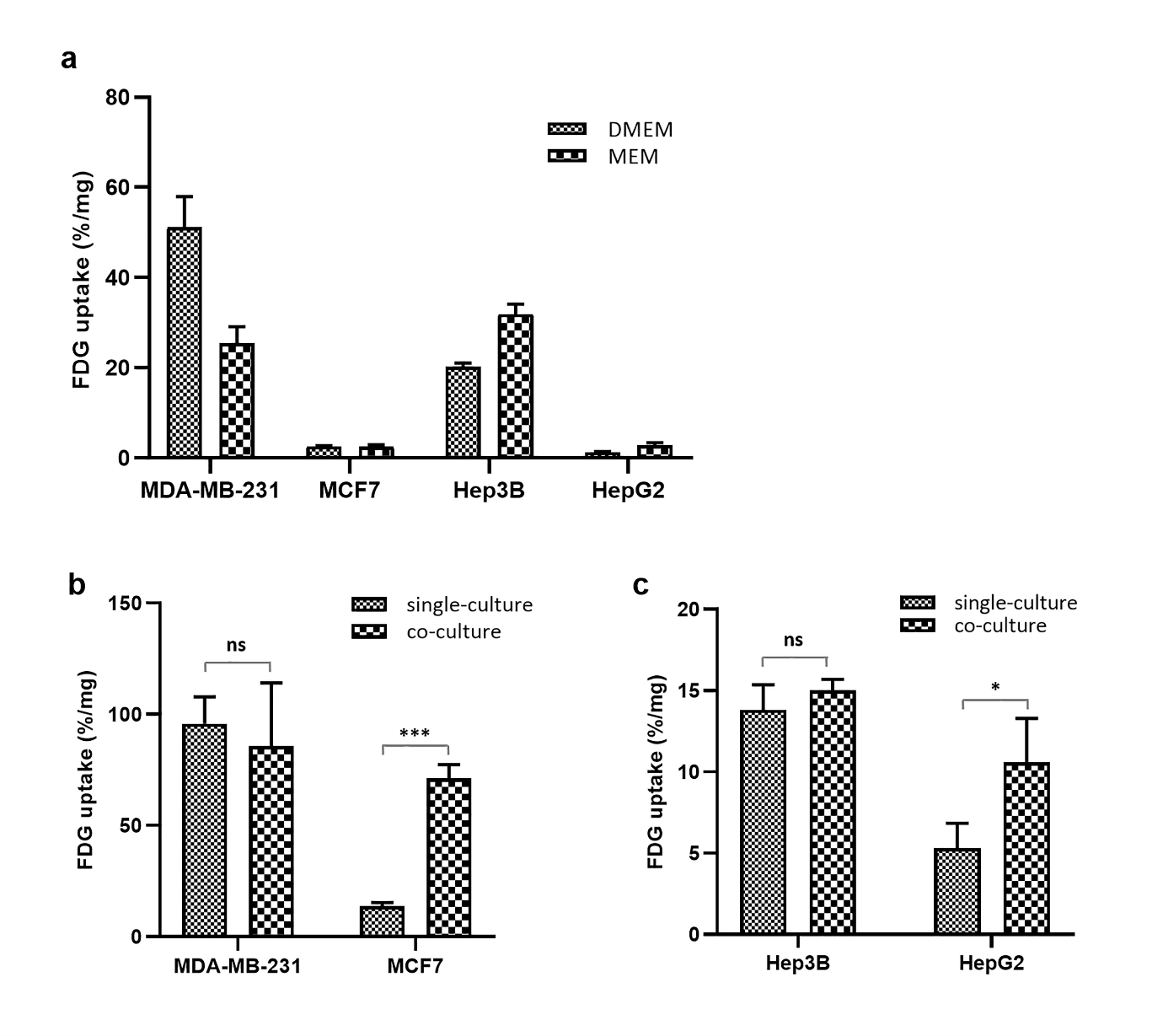

**Supplementary Figure S2**

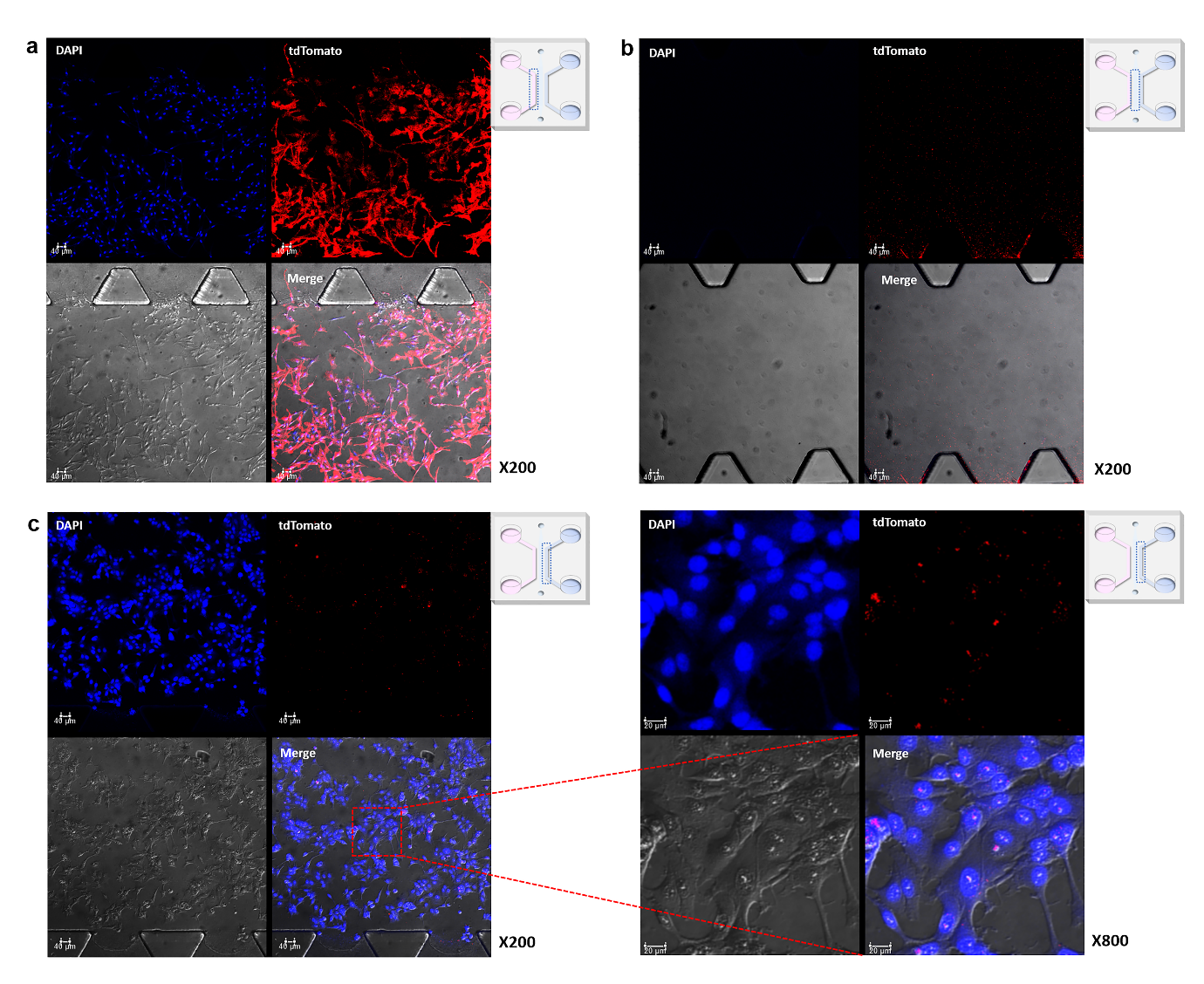

**Supplementary Figure S3**

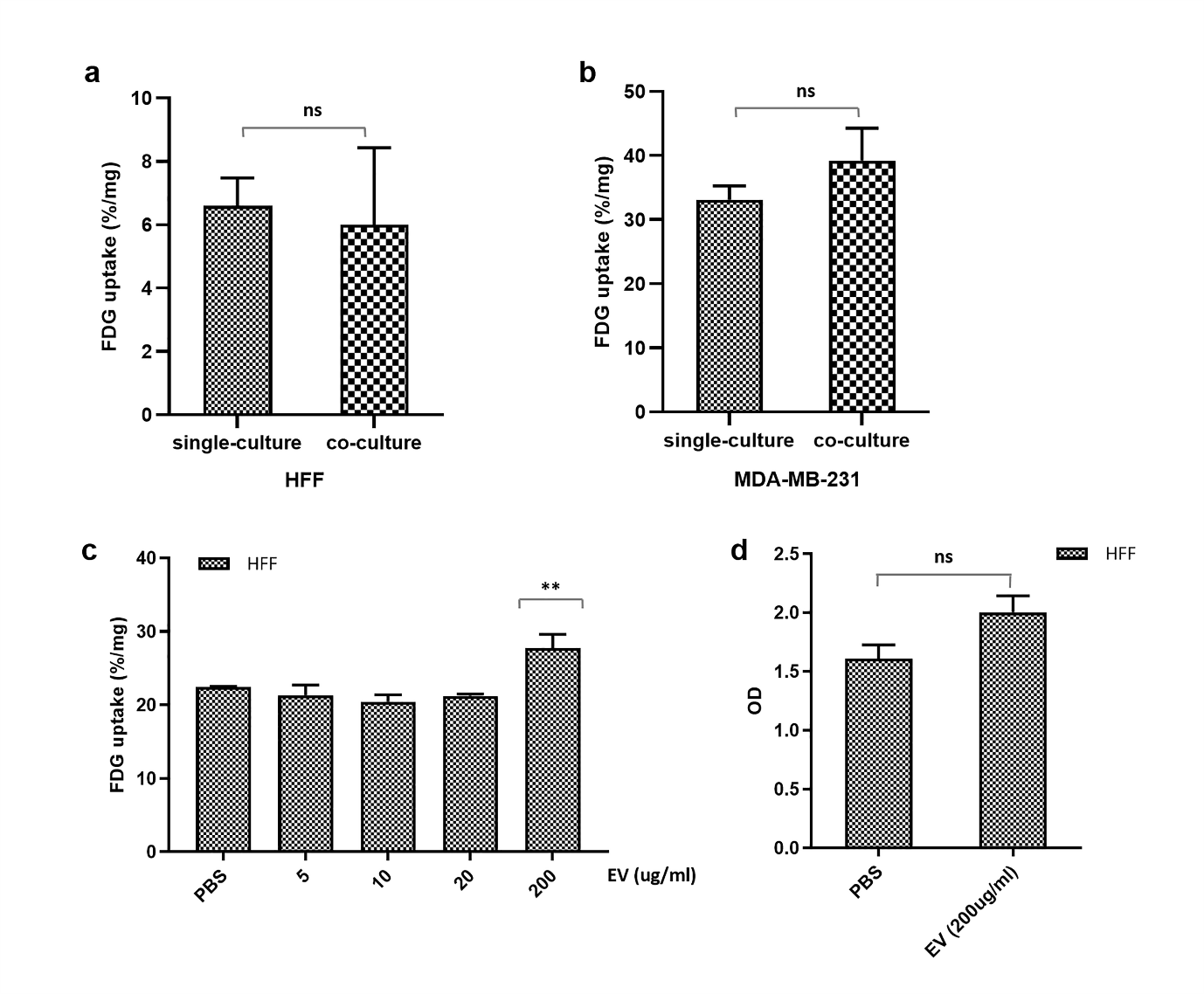

**Supplementary Figure S4**

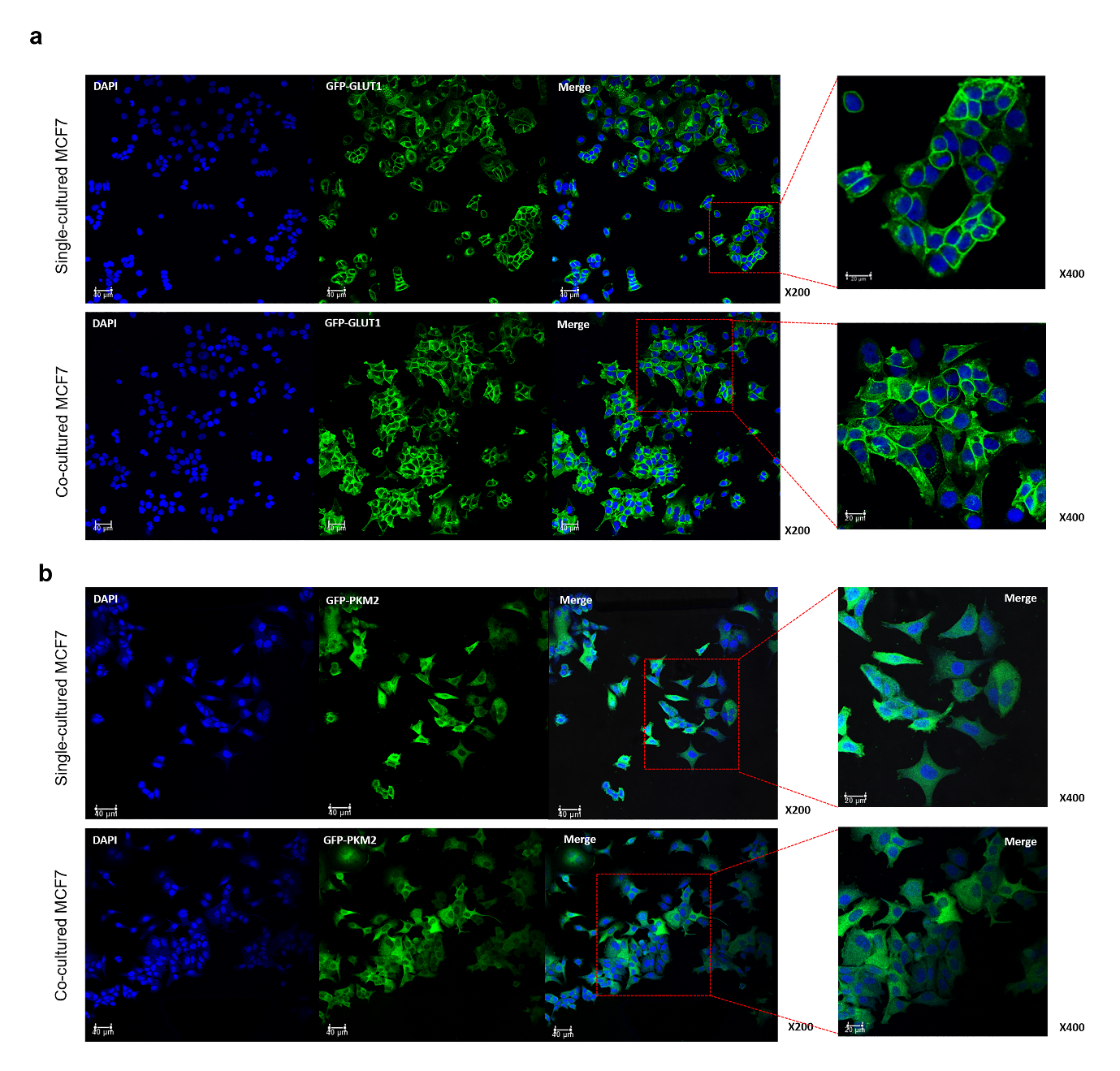

**Supplementary Figure S5**

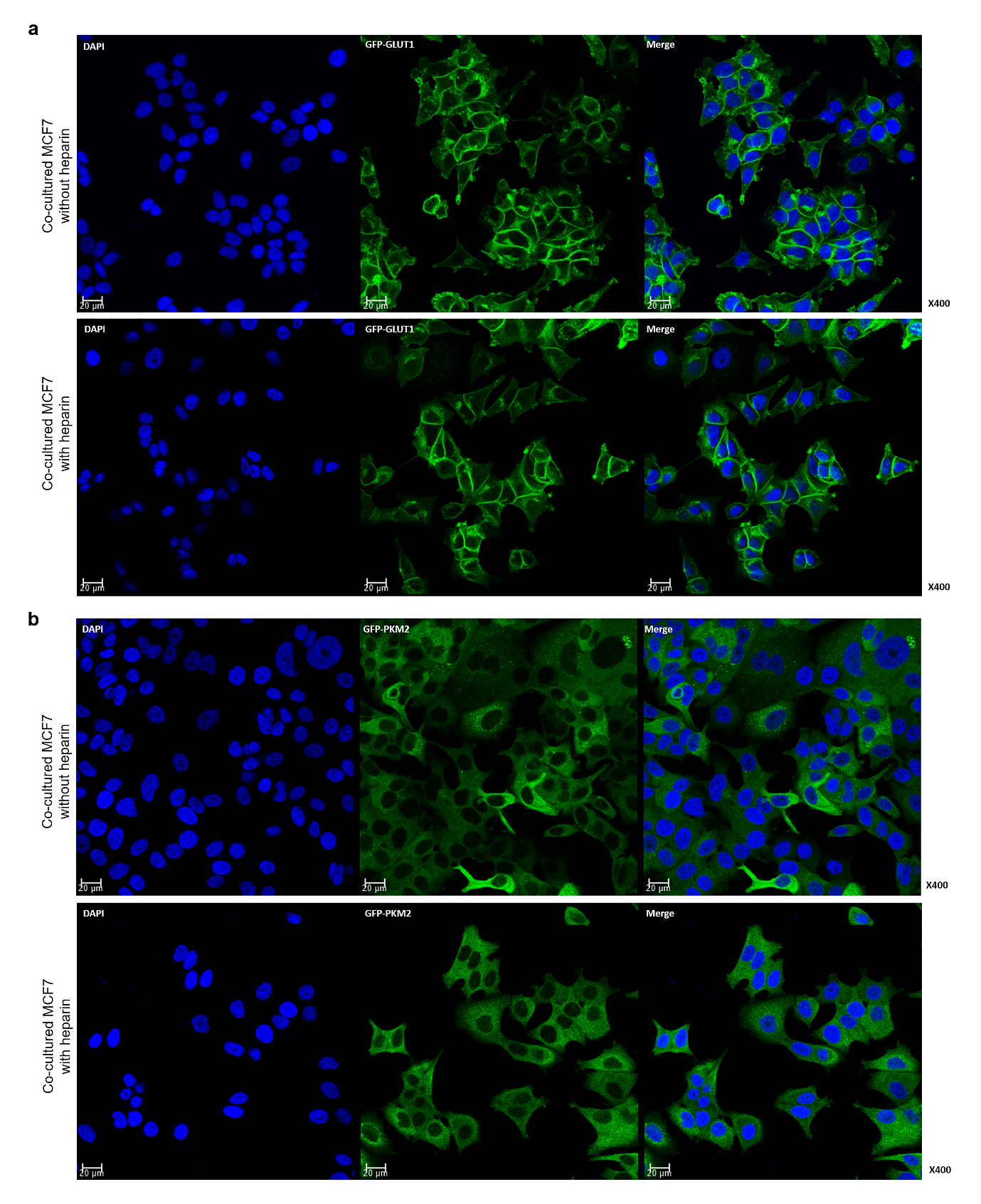

**Supplementary Figure S6**

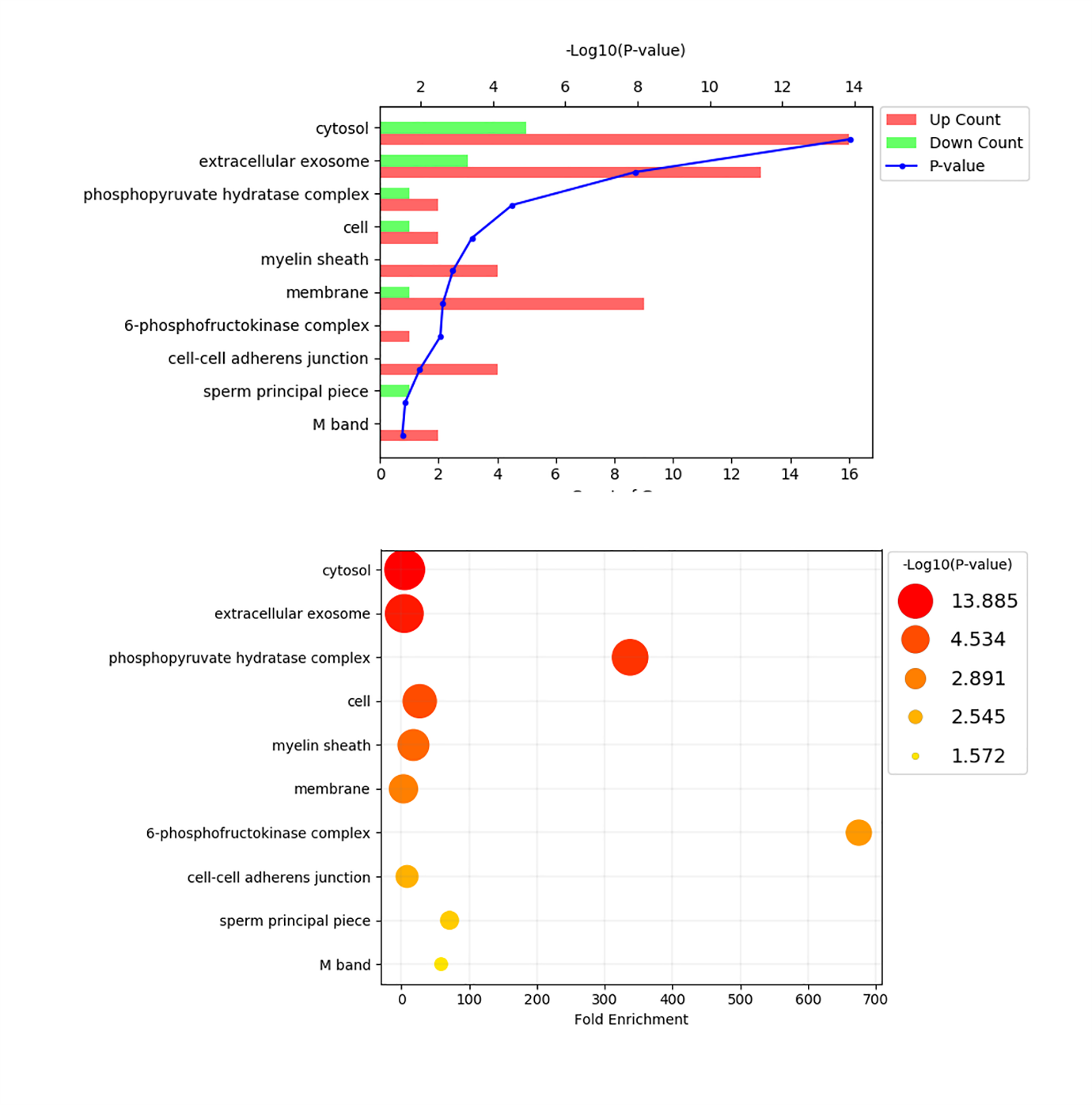

**Supplementary Figure S7**

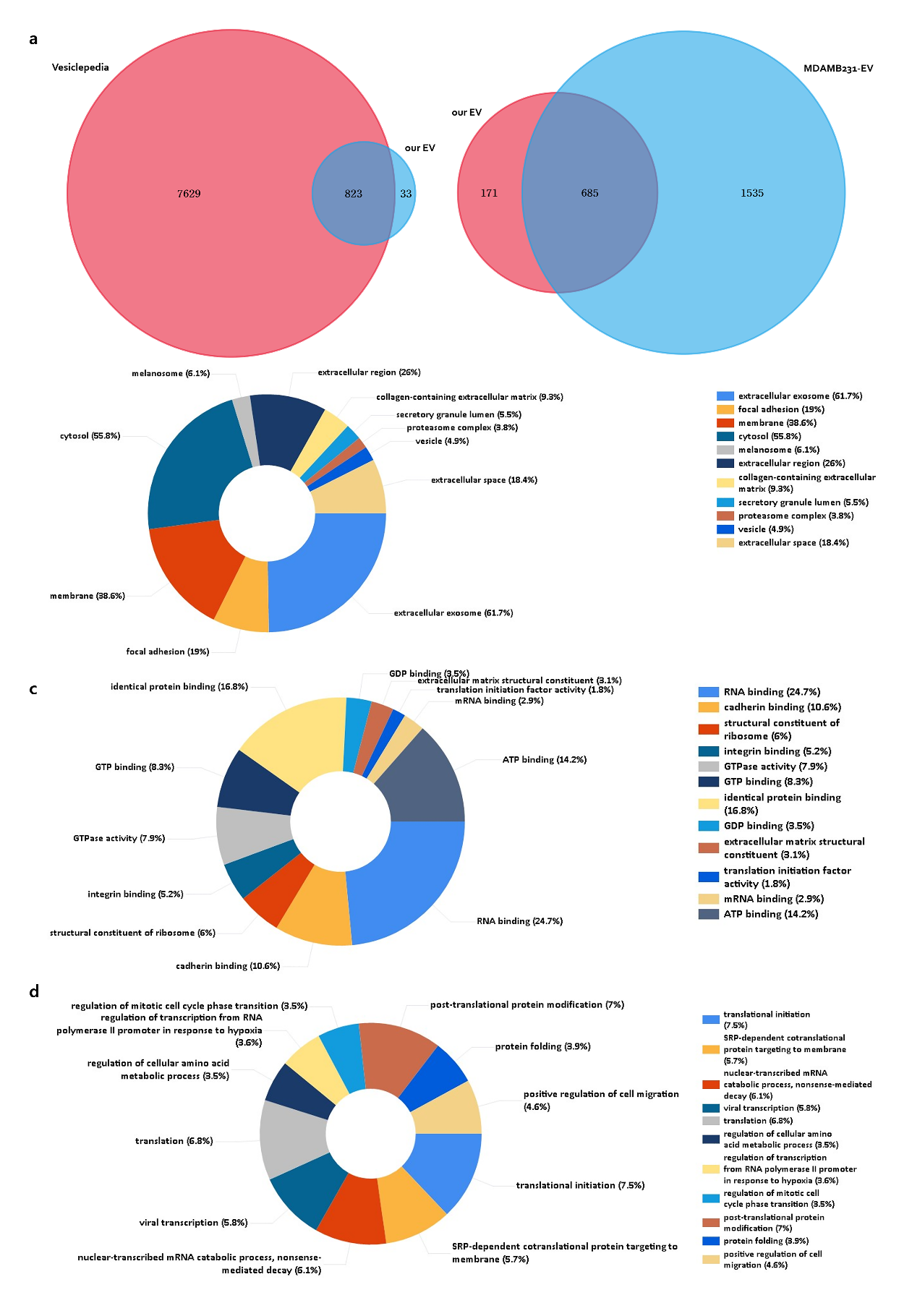

**Supplementary Figure S8**

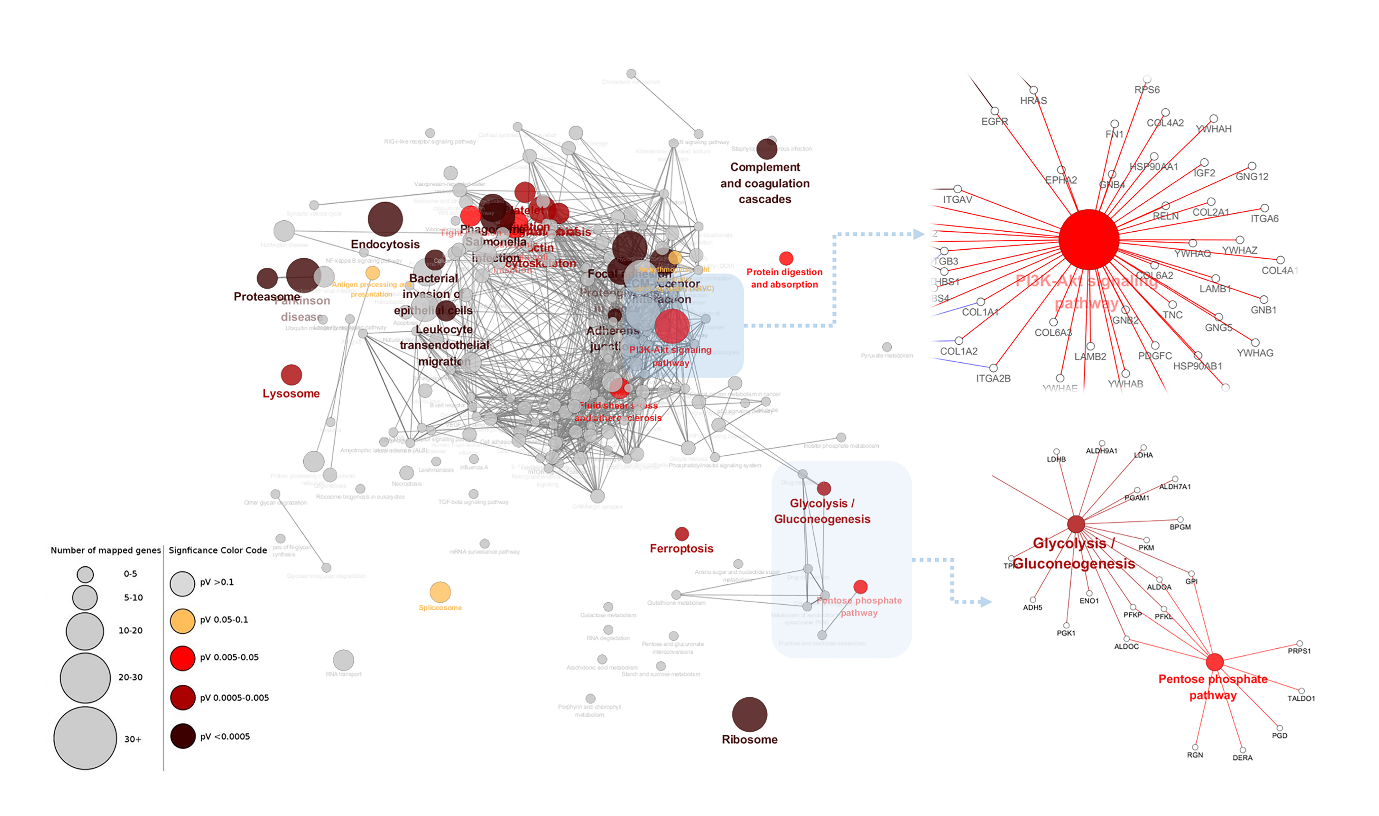

**Supplementary Figure S9**

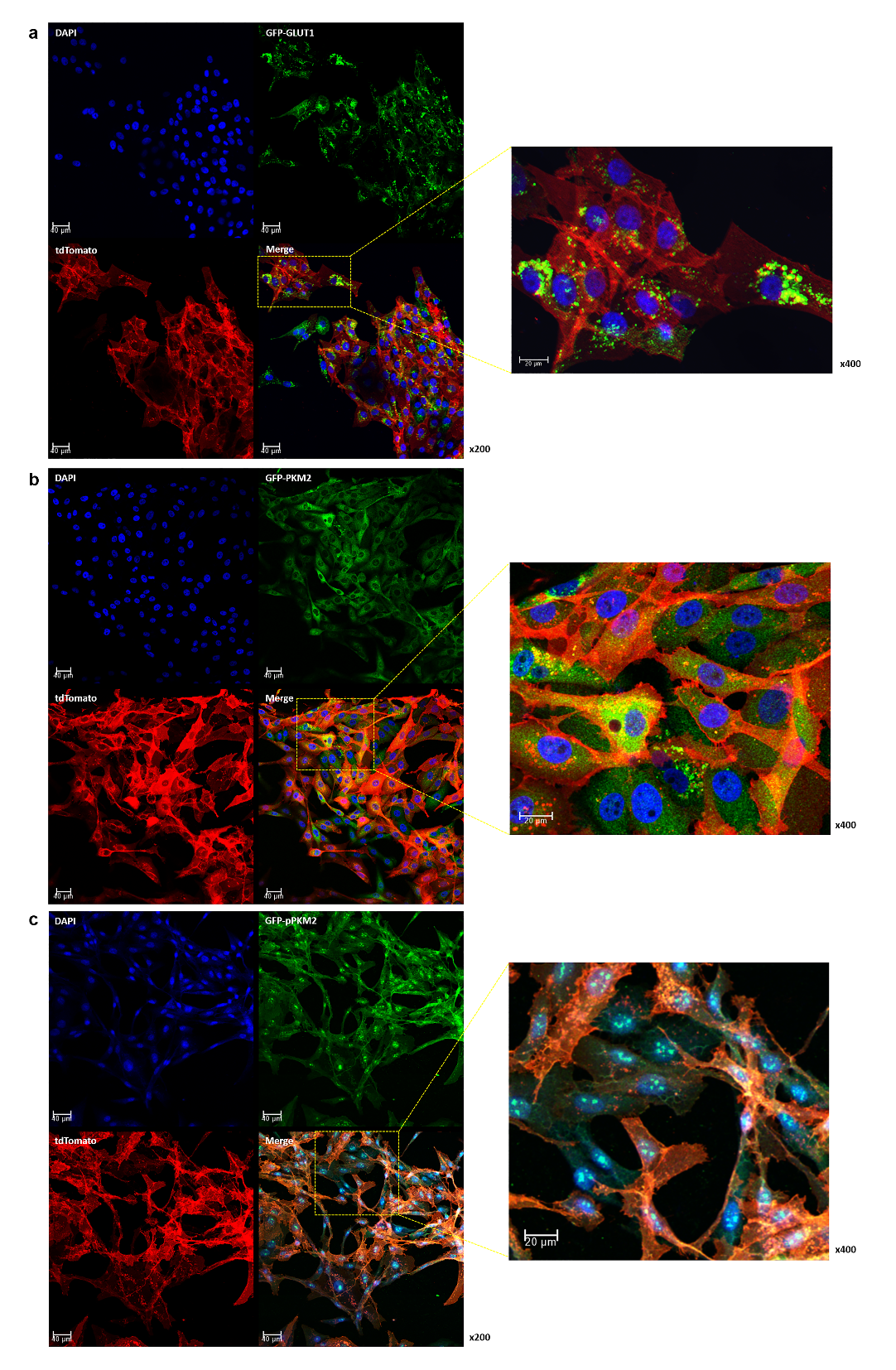

**Supplementary Figure Captions**

**Supplementary Figure S1**. Evaluation of FDG uptake impact under various conditions; Comparison of the effect of glucose amount in the medium on FDG uptake (a); No significant change of FDG uptake in MDA-MB-231 cells after co-culture with MCF7 cells (b); Changes in FDG uptake by co-culture between Hep3B and HepG2 (c).

**Supplementary Figure S2**. Confocal microscopic images of MDA-MB-231-tdTomato cells and tdTomato-EVs in the donor (a), middle (b) and recipient channels (c) of microfluidic chip.

**Supplementary Figure S3**. Change of FDG uptake in the HFF cells. Change of FDG uptake in HFF cells (a) co-cultured with MDA-MB-231 cells (b); MDA-MB-231-derived EVs have significant effect of FDG uptake in HFF cells (c); MDA-MB-231-derived EVs did not activate proliferation in HFF cells (d) unlike in MCF7 cells.

**Supplementary Figure S4**. Confocal microscopic images showing expression level of PKM2 and GLUT1 in MCF7 cells. Visually, there were no clear changes in GLUT1 (a) and PKM2 (b) expression in MCF7 cells after co-culture with MDA-MB-231 cells on confocal microscopic images.

**Supplementary Figure S5**. Confocal microscopic images showing expression level of PKM2 and GLUT1 in MCF7 cells after inhibition of EV uptake using heparin. Visually, there were no significant changes in GLUT1 (a) and PKM2 (b) expression in MCF7 cells after heparin treatment. However, cell proliferation has been inhibited after heparin treatment.

**Supplementary Figure S6**. The extracellular exosome was the second most cellular component after the cytosol in the analysis of the cellular components of genes related to glycolysis in the entire transcriptomic data.

**Supplementary Figure S7**. Venn diagram describing the matched proteins with total EV and MDA-MB-231 EV database from Vesiclopedia (a). The GO analysis of the 856 proteins identified in the MDA-MB-231-derived EVs; (b) cellular component, (c) molecular function, and (d) Biologic process

**Supplementary Figure S8**. KEGG pathway analysis of MDA-MB-231-derived EVs. Among the significant KEGG pathways, glycolysis/glucogenes and PI3K-Akt signal pathways were identified to confirm the presence of important proteins responsible for glycolysis and PKM2 phosphorylation inside EVs.

**Supplementary Figure S9.** The expression pattern of GLUT1 and PKM2 in MDA-MB-231 cells. Most of the GLUT1 proteins was located in the cytosol like granules, which may be highly expressed GLUT1 proteins located in the endoplasmic reticulum or Golgi apparatus (a). Expression (b) and (c) phosphorylation of PKM2 in MDA-MB-231 cells was highly activated.
